# The lipid droplet protein DHRS3 is a regulator of melanoma cell state

**DOI:** 10.1101/2024.03.25.586589

**Authors:** Eleanor Johns, Yilun Ma, Pakavarin Louphrasitthipol, Christopher Peralta, Miranda V. Hunter, Jeremy H. Raymond, Henrik Molina, Colin R. Goding, Richard M. White

**Author notes:** Correspondence to Richard M. White.

## Abstract

Lipid droplets are fat storage organelles composed of a protein envelope and lipid rich core. Regulation of this protein envelope underlies differential lipid droplet formation and function. In melanoma, lipid droplet formation has been linked to tumor progression and metastasis, but it is unknown whether lipid droplet proteins play a role. To address this, we performed proteomic analysis of the lipid droplet envelope in melanoma. We found that lipid droplet proteins were differentially enriched in distinct melanoma states; from melanocytic to undifferentiated. DHRS3, which converts all-trans-retinal to all-trans-retinol, is upregulated in the MITF^LO^/undifferentiated/neural crest-like melanoma cell state and reduced in the MITF^HI^/melanocytic state. Increased DHRS3 expression is sufficient to drive MITF^HI^/melanocytic cells to a more undifferentiated/invasive state. These changes are due to retinoic acid mediated regulation of melanocytic genes. Our data demonstrate that melanoma cell state can be regulated by expression of lipid droplet proteins which affect downstream retinoid signaling.

## Introduction

Cell plasticity is one of the defining features of melanoma, with significant differences in clinical outcomes associated with phenotype switching.^1–3^ Melanoma cells can reversibly move between different transcriptional states, each of which is associated with distinct phenotypes. These different cell states can have important implications in disease progression, metastasis, and drug resistance.^2,4–6^ Numerous studies have attempted to define transcriptomic signatures that identify these diverse melanoma cell states, which have begun to converge on at least 4 distinct classifications: undifferentiated, neural crest-like, transitory, and melanocytic.^3–5,7^ What drives each and what allows cells to move between states, remain significant open questions.

Across a multitude of tumor types, uptake of fatty acids has been shown to be supportive of a variety of tumor metabolic needs which can facilitate tumor formation, cancer cell survival, and metastasis.^8–11^ Lipid droplets, which are associated with excess fatty acid levels, have long been studied for their roles in fatty acid and lipid storage, particularly in specialized cells such as adipocytes. However, there is increasing interest in understanding the complex role of lipid droplets in cellular metabolism across a diversity of cell types and disease states. Recent research has clearly demonstrated the importance of lipid droplet formation and function in melanoma,^12–14^ however we do not fully understand how lipid droplet formation affects disease progression. Furthermore, the dynamic regulation of the lipid droplet in this lineage remains an open area of investigation.

Previous work from our laboratory has demonstrated that formation of lipid droplets is associated with changes in cell state. During melanoma development, tumor cells take up diverse species of fatty acids from nearby adipocytes and readily form lipid droplets.^14^ Uptake of saturated fatty acids is linked to a more undifferentiated, invasive cell state.^15^ Conversely, melanocytic melanoma cells, which are more oxidative, rely on increased lipid droplet content to support unsaturated fatty acid flux through the mitochondria.^12^ These observations imply that the regulation and function of lipid droplets is highly dependent on cellular context and substrate availability. Furthermore, they suggest that lipid droplets may be linked to distinct transcriptional cell states.

While significant work has focused on the lipid components of the droplet, many studies also focus on the protein coat which surrounds the lipid rich core.^16^ Differential localization of proteins to the lipid droplet membrane can have dramatic effects on lipid droplet function as well as cellular metabolism.^17–19^ Numerous proteomic studies have been conducted to better understand the role of these lipid droplet proteins in a variety of contexts.^19–23^ These observations led us to hypothesize that proteins in the lipid droplet coat might be determinants of melanoma cell state.

To better understand the potential role of lipid droplets in determining melanoma phenotypic identity we used APEX2 proximity labeling^20,24^ to define the melanoma lipid droplet proteome. Using a combination of transcriptomic and biochemical approaches, we demonstrate that a particular lipid droplet protein, DHRS3, plays a critical role in supporting the undifferentiated/neural crest-like melanoma cell state.

## Results

### Proximity labeling of the melanoma lipid droplet proteome

We hypothesized that a key determinant of the effect of lipid droplet accumulation in melanoma would be due to the dynamic composition of the lipid droplet protein coat. As no previous study had examined the melanoma lipid droplet proteome, we adapted lipid droplet targeted APEX2 proximity labeling^20,24^ for our cell type of interest. This approach utilizes a lipid droplet specific APEX2 (by fusion to the lipid droplet protein Perilipin 2 (PLIN2). We introduced an inducible lipid droplet targeted PLIN2-V5-APEX2 construct along with a cytosolic, Cyto-V5-APEX2 control to A375 human melanoma cells. (Figure 1A). Titration of doxycycline for 48 hours showed dose responsive expression of the APEX2 fusion constructs (Figure 1B). In addition, treatment with biotin phenol and H_2_O_2_ demonstrated robust, inducible biotinylation of proteins in both conditions (Figure 1B, Supp Fig 1C). To collect the large numbers of lipid droplets necessary for proteomics,^20^ we treated cells with 200 μM Oleic Acid for 24 hours (Supp Fig 1A-B). Imaging of the PLIN2-V5-APEX2 and Cyto-V5-APEX2 cell lines showed clear localization of APEX2 to the lipid droplet combined with an enrichment of biotinylated proteins around the lipid droplet in PLIN2-V5-APEX2 cells only (Figure 1C).

**Figure 1.**
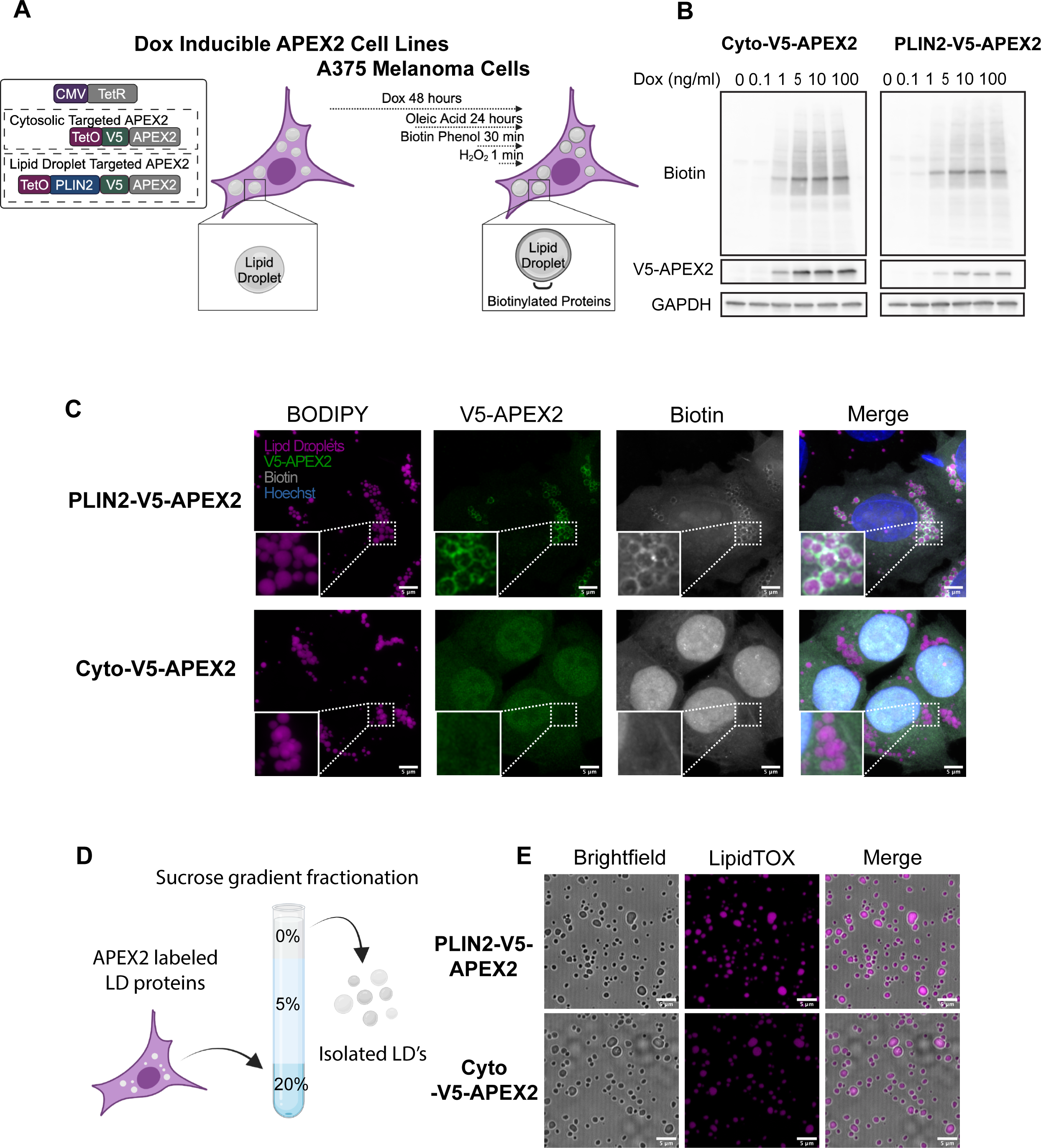
Proximity labeling of the melanoma lipid droplet proteome. A. Schematic of the APEX2 expressing cell lines used in this study B. Western blot of cells treated with indicated doses of doxycycline for (48 hours) and 200 μM Oleic Acid (24 hours) followed by 30 minutes of biotin phenol (500 μM) and 1 minute of H_2_O_2_ (1 mM). C. Immunofluorescence imaging of APEX2 expressing cells treated as in B followed by fixation and staining for lipid droplets (BODIPY), PLIN2-v5-APEX2 (V5 antibody) and biotin (Streptavidin antibody). Representative high-resolution images from n = 18 images taken from n = 3 independent experiments. Scale bars are 5 μm. D. Simplified schematic of gradient ultracentrifugation to isolate lipid droplets (see methods). E. Confocal imaging of lipid droplets stained with the lipid droplet dye LipidTOX after isolation. All data are representative of at least n = 3 independent experiments. Scale bars are 5 μm. Diagrams made with Biorender.com

Given that we observed labeling of cytosolic proteins in PLIN2-V5-APEX2 cells, as has been previously reported,^20^ we performed a further enrichment of lipid droplets by combining the proximity labeling approach with gradient ultracentrifugation (Figure 1D).^20^ Using this method, we were able to isolate intact lipid droplets relatively devoid of common contaminants (Figure 1E, Supp Fig 1D).

### Identification of the melanoma lipid droplet proteome

To characterize the lipid droplet proteome in melanoma cells we performed mass spectrometry. We induced APEX2 activity followed by gradient ultracentrifugation to isolate biotinylated lipid droplet fractions. Western blotting demonstrated effective isolation of a biotinylated fraction in PLIN2-V5-APEX2 cells treated with H_2_O_2_ (Figure 2A), but not in Cyto-V5-APEX2 (Figure 2B) or no H_2_O_2_ controls (Figure 2C). After isolation of lipid droplet fractions, we used streptavidin immunoprecipitation to enrich for the biotinylated proteins followed by LC-MS/MS (Figure 2D). Analysis revealed clear differentiation between samples depending on condition (Supp Figure 2) and enrichment of canonical lipid droplet proteins in the PLIN2-V5-APEX2 condition only (Supp Figure 3A-B).

**Figure 2.**
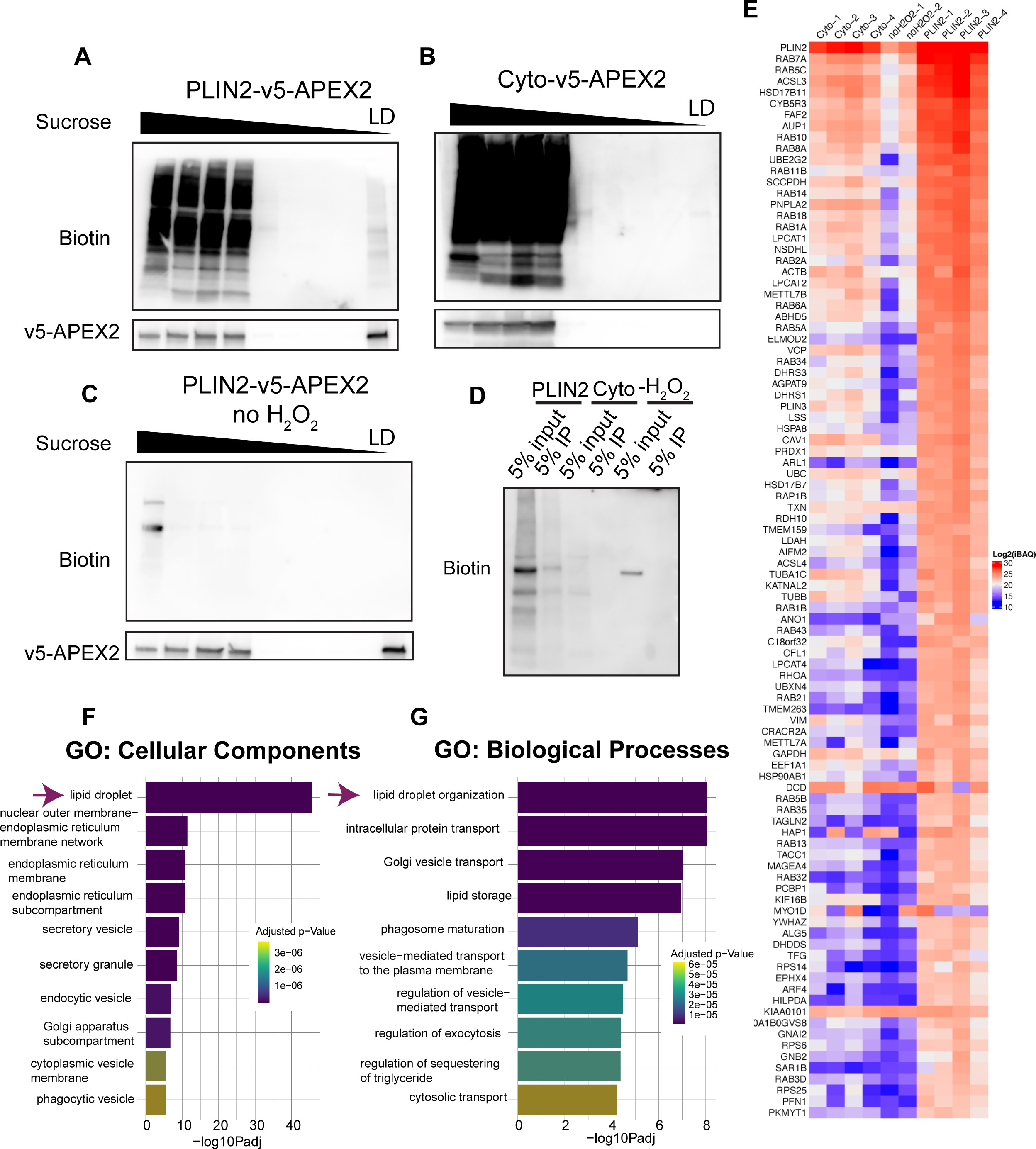
Identification of the melanoma lipid droplet proteome. A-C. Representative western blots (from n = 4 independent experiments) of equal percent by volume of each fraction after APEX2 labeling and gradient ultracentrifugation as described in Methods. D. Representative western blot of streptavidin immunoprecipitation of isolated lipid droplet proteins from the indicated conditions which were used for LC-MS/MS E. Heatmap of the log_2_iBAQ values for the top 96 lipid droplet proteins identified by LC-MS/MS. F-G. Top 10 pathways as identified using gProfiler. Data are representative of 4 independent experiments (Cyto and PLIN2-APEX2) and 2 independent experiments for -H_2_O_2_.

**Figure 3.**
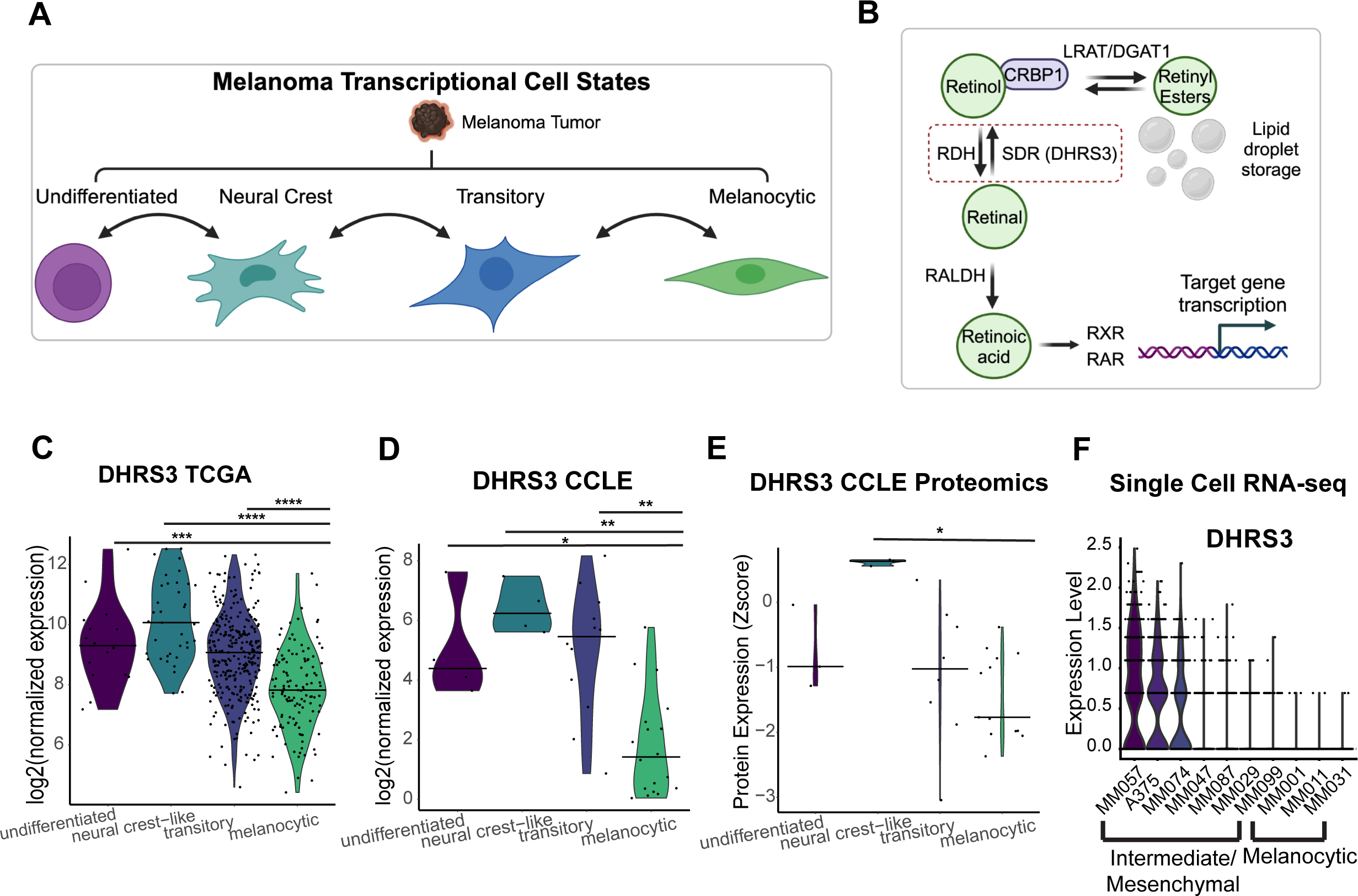
DHRS3 is a lipid droplet protein enriched in the undifferentiated versus melanocytic cell state. A. Diagram of the four melanoma transcriptional cell states as in.^4,82^ B. Simplified diagram of retinoic acid metabolism, highlighting the known function of DHRS3. C-D. Violin plots of analysis of the CCLE and TCGA transcriptomic databases (bar = median, dots represent each sample or cell line.). Data was grouped by the Tsoi cell state classifiers which have been previously annotated for this data.^4^ Statistics were generated with two-sided Wilcoxon rank sum test with Holm correction. * p < 0.05 **, p < 0.01, *** p < 0.001, **** p < 0.0001. E. Analysis of the CCLE proteomics database. Data was grouped by Tsoi cell state classifiers. Statistics were generated with two-sided Wilcoxon rank sum test with Holm correction. * p < 0.05. F. mRNA expression of DHRS3 in scRNA-seq of patient-derived melanoma cell lines from Wouters et al., 2020. Diagrams made with Biorender.com

To generate a list of lipid droplet proteins, we used label free quantification based on the Intensity Based Absolute Quantification (iBAQ) value for each protein. To account for the cytosolic background and nonspecific binding to the beads we subtracted the average Cyto-V5-APEX2 iBAQ values from the PLIN2-V5-APEX2 iBAQ values to generate a normalized iBAQ value (Supplemental Table 2).^24^ To determine a high confidence list of lipid droplet proteins, we set a cut-off such that 60% of previously validated lipid droplet proteins were identified^20^ (Supplemental Table 1), which yielded a total of 96 proteins (Figure 2E, Supplemental Table 3). Analysis of this list with gProfiler demonstrated enrichment of lipid droplet genes as the top pathway (Figure 2F-G). We were therefore able to determine a robust list of 96 lipid droplet proteins in melanoma.

### DHRS3 is a lipid droplet protein enriched in the undifferentiated versus melanocytic cell state

We next sought to better understand the function of these lipid droplet proteins in melanoma. We were most interested in lipid droplet proteins that might have a function in specific melanoma phenotypes, as opposed to general functions that occur in most cell types. Because it is now well-recognized that human melanomas exist in at least 4 distinct transcriptional states, we asked whether any of the identified proteins were specifically enriched in certain states. To do this, we compared our proteomics data to published melanoma transcriptional cell state signatures which have been linked to these phenotypic states: undifferentiated, neural crest-like, transitory, and melanocytic (Figure 3A).^4^ We decided to focus on one protein, *DHRS3*, for further downstream functional analysis. *DHRS3* is the enzyme that converts all-trans-retinal to all-trans-retinol^25,26^ (Figure 3B, Supp Fig 4A). This was especially intriguing to us given the known role of retinoid signaling in melanocyte differentiation^27,28^ as well as previous data which identified Retinoid X Receptors (RXRs) as a therapeutic vulnerability of the neural crest-like melanoma cell state.^2^ We found that *DHRS3* was markedly enriched in the undifferentiated and neural crest-like melanoma cells (Figure 3C-D) compared to the melanocytic/pigmented state, where it is minimally expressed. This made it a good candidate for studying its functional role in these very different phenotypic cell states. As we generally see similar trends in DHRS3 levels between the undifferentiated and neural crest-like states vs. the melanocytic state, we will be referring to this less differentiated group as undifferentiated/neural crest-like for the rest of the paper. Confirming the link between upregulation of *DHRS3* mRNA and protein, DHRS3 protein was also upregulated in neural crest-like cells in the CCLE proteomics dataset (Figure 3E).^29^ We also analyzed a previously published human melanoma single cell RNA-seq dataset^5^ composed of expression data from patient-derived cell lines classified as either ‘mesenchymal’, ‘intermediate’ or ‘melanocytic’. Similar to what we observed in the TCGA and CCLE bulk RNA-seq datasets, *DHRS3* was highly expressed in mesenchymal/intermediate-like cell lines and downregulated in melanocytic cell lines (Figure 3F). Taken together, our data suggests that DHRS3 mRNA and protein expression is associated with the undifferentiated/neural crest-like cell state in melanoma, raising the question of whether it plays a functional role.

**Figure 4.**
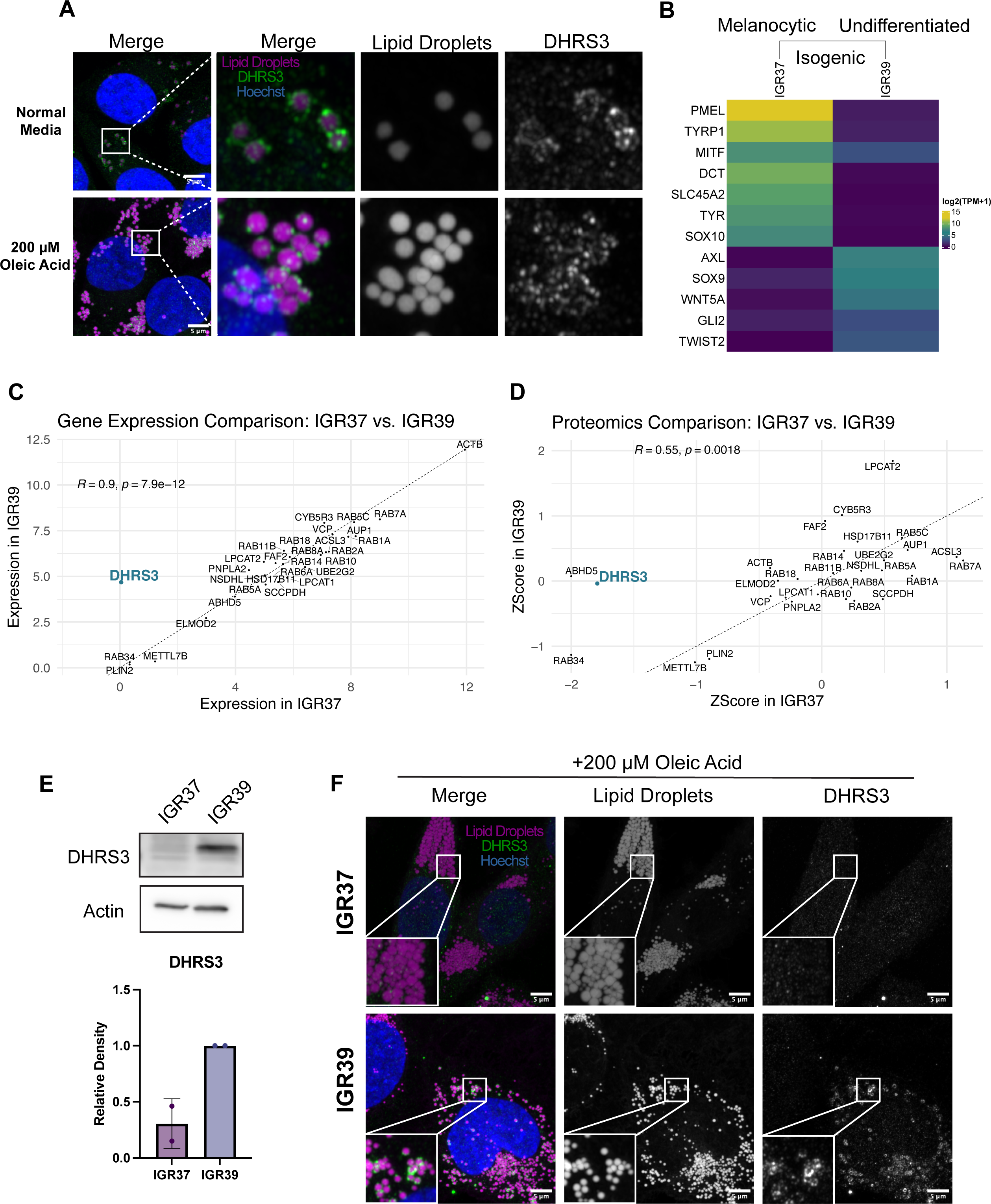
DHRS3 is enriched in undifferentiated melanoma cells. A. Representative images of staining for endogenous DHRS3 and lipid droplets (BODIPY) in A375 cells. Where indicated cells were treated with oleic acid for 24 hours. Images are representative of 2 independent experiments. Scale bars are 5 μm. B. Heatmap of log2 Normalized Counts (from the CCLE) of some representative genes known to differentiate undifferentiated vs. melanocytic cells. C. Scatter plot comparing the expression level (log2 Normalized Counts) of the top 30 lipid droplet genes in the melanoma lipid droplet proteome in IGR37 vs. IGR39 melanoma cells. DHRS3 is highlighted on the graph. Correlation calculated with Spearman Correlation Coefficient. D. Scatter plot comparing the protein level (ZScore) of the top 30 lipid droplet proteins in the melanoma lipid droplet proteome in IGR37 vs. IGR39 cells. DHRS3 is highlighted in the graph. Correlation calculated with Spearman Correlation Coefficient. E. Western blot with quantification of DHRS3 protein levels. Representative of 2 independent experiments. Data are presented as mean +/− SD. F. Representative images of IGR37 and IGR39 cells treated with oleic acid for 24 hours. Cells were fixed & stained with LipidTOX and an antibody against endogenous DHRS3. Data from 3 independent experiments. Scale bars are 5 μm.

### DHRS3 is an effector of melanoma cell state

DHRS3 has been previously identified in lipid droplets,^19,30,31^ but its functional role remains incompletely understood, particularly in the context of melanoma. We first confirmed our proteomic results by immunofluorescence. Consistent with our data, we found that DHRS3 exclusively localizes to the lipid droplet in A375 cells in both normal media and media with oleic acid (Figure 4A).

To study the functional relationship between DHRS3 expression and melanoma cell state, we sought to use an isogenic cell line system that represented these different states. A unique resource in melanoma are the IGR37/IGR39 cell lines. These are two cell lines that come from the same melanoma patient but have markedly different characteristics. The IGR37 line is melanocytic, with high levels of *MITF* which readily pigments. The IGR39 line is undifferentiated, does not express *MITF*, and is unpigmented. Furthermore, IGR37 cells are sensitive to BRAF inhibition while IGR39 cells are resistant and invasive.^32^ This pair of lines best represents the melanocytic and undifferentiated cell states respectively ^4^ (Figure 4B). At both the RNA and protein level, DHRS3 is markedly higher in the IGR39 (MITF^LO^, undifferentiated) cells compared to the IGR37 (MITF^HI^, melanocytic) cells (Figure 4C-E), consistent with our TCGA/CCLE analysis above. Moreover, when we compared the top 30 proteins we found at the lipid droplet by proteomics, DHRS3 is the sole protein that reliably distinguishes the IGR37 vs. IGR39 cell line (Figure 4C-D). Consistent with this, when we treated cells with oleic acid, DHRS3 was enriched at the lipid droplet in IGR39 cells but not IGR37 cells (Figure 4F). To extend this observation to other cell lines, we performed imaging of additional undifferentiated/neural crest-like cell lines, which revealed a similar localization of DHRS3 to the lipid droplet (Supp Figure 4B-C).

To test the role of DHRS3 in melanoma cell state, we overexpressed DHRS3 in the IGR37 (melanocytic) cells, which normally have very low levels of the protein (Figure 5A-B). Consistent with the hypothesis that DHRS3 promotes de-differentiation of melanoma cells, overexpression of DHRS3 in IGR37 cells resulted in a visible decrease in pigmentation (Figure 5C). To define the downstream targets of DHRS3 in melanoma, we performed bulk RNA-sequencing on IGR37 DHRS3 overexpressing versus control cells. This revealed a total of 297 differentially expressed genes (Supp. Figure 5A, Supplementary Table 4-5). To determine whether overexpression of DHRS3 induced transcriptional changes in IGR37 cells representative of de-differentiation, we used Gene Set Enrichment Analysis (GSEA) to calculate an enrichment score for each of the previously published gene sets corresponding to melanoma cell state.^4^ Cells overexpressing DHRS3 highly upregulated genes corresponding to the undifferentiated/neural crest-like states (Figure 5D, Supp Figure 5B-C). GSEA also revealed that pathways related to tumor invasion and epithelial-to-mesenchymal transition were highly enriched in DHRS3^OE^ cells (Figure 5E), consistent with a less differentiated cell state.^3,4,33^

**Figure 5.**
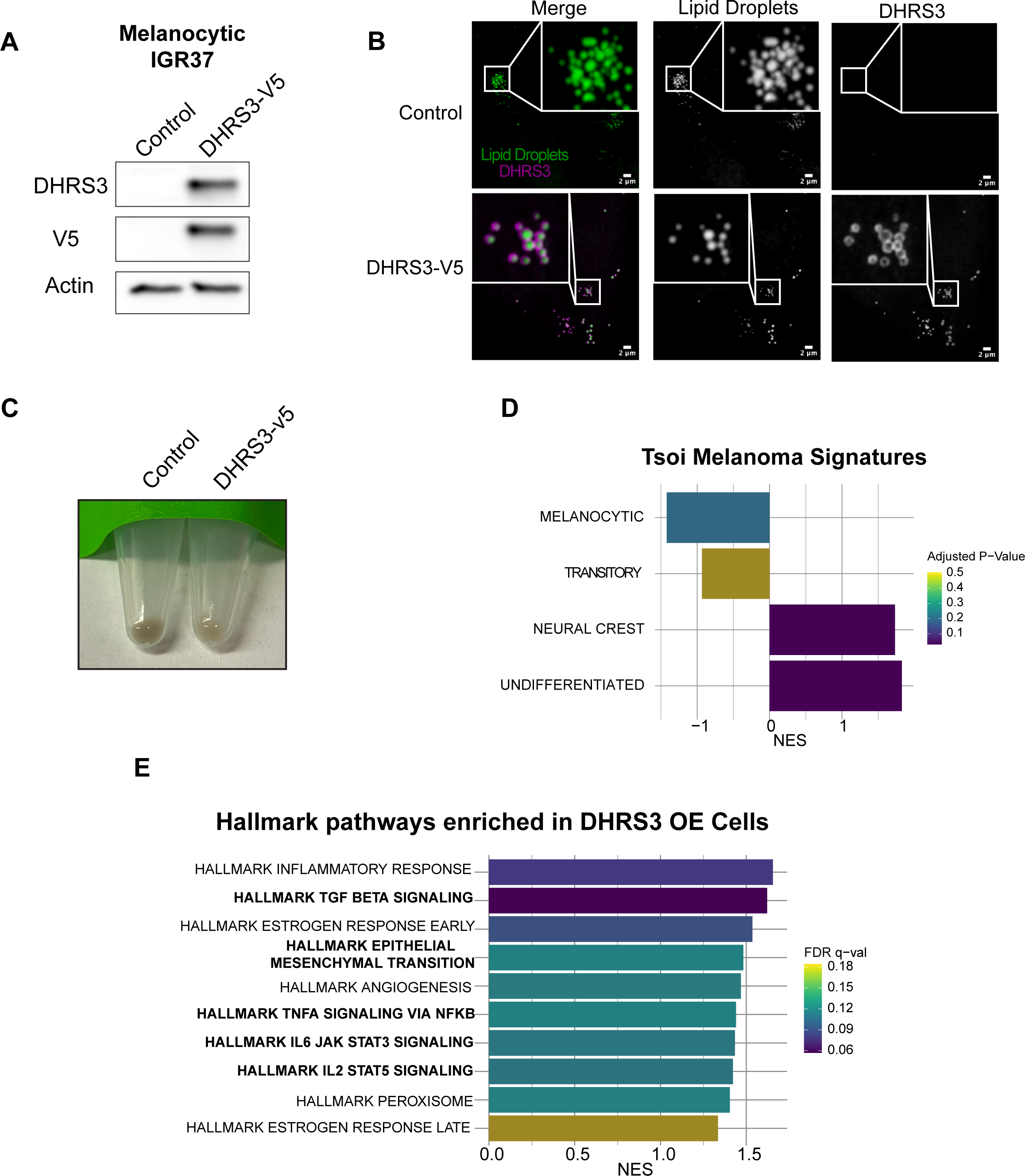
DHRS3 is an effector of melanoma cell state. A. Representative western blot of DHRS3 overexpression. Data from 3 independent experiments. B. Representative images of DHRS3 overexpressing and control IGR37 cells. Data from n = 2 independent experiments. Scale bars are 2 μm. C. Representative image of pellet color in Control and DHRS3 overexpressing cells. Cells were counted prior to spinning down to ensure equal numbers. Data representative of at least 3 independent experiments. D. GSEA results using log2FC generated by DESeq2 using custom pathways generated from the Tsoi classifications ^4^. E. GSEA results using normalized counts from bulk RNA-seq. Hallmark pathways related to invasion and EMT are shown in bold. RNA-seq data was collected in triplicate.

### DHRS3 regulates differentiation through retinoid signaling

We next performed a broader pathway analysis using the Curated Gene Sets from MSigDB. Amongst the enriched pathways were those related to retinoid metabolism, which is in line with the known function of DHRS3^25^ (Figure 6A). In further support of a relationship between DHRS3 overexpression and retinoid metabolism, when we performed motif analysis to identify conserved transcription factor binding motifs within the promoter regions of genes upregulated upon DHRS3^OE^, we found significant enrichment of a HOX motif (Supp Figure 5D). HOX genes are well established downstream targets of RXR/RAR signaling.^34,35^. Enrichment of the HOX motif further suggested to us that RXR/RAR activity was impacted, which in turn caused changes in HOX gene activity and regulation of downstream genes.

**Figure 6.**
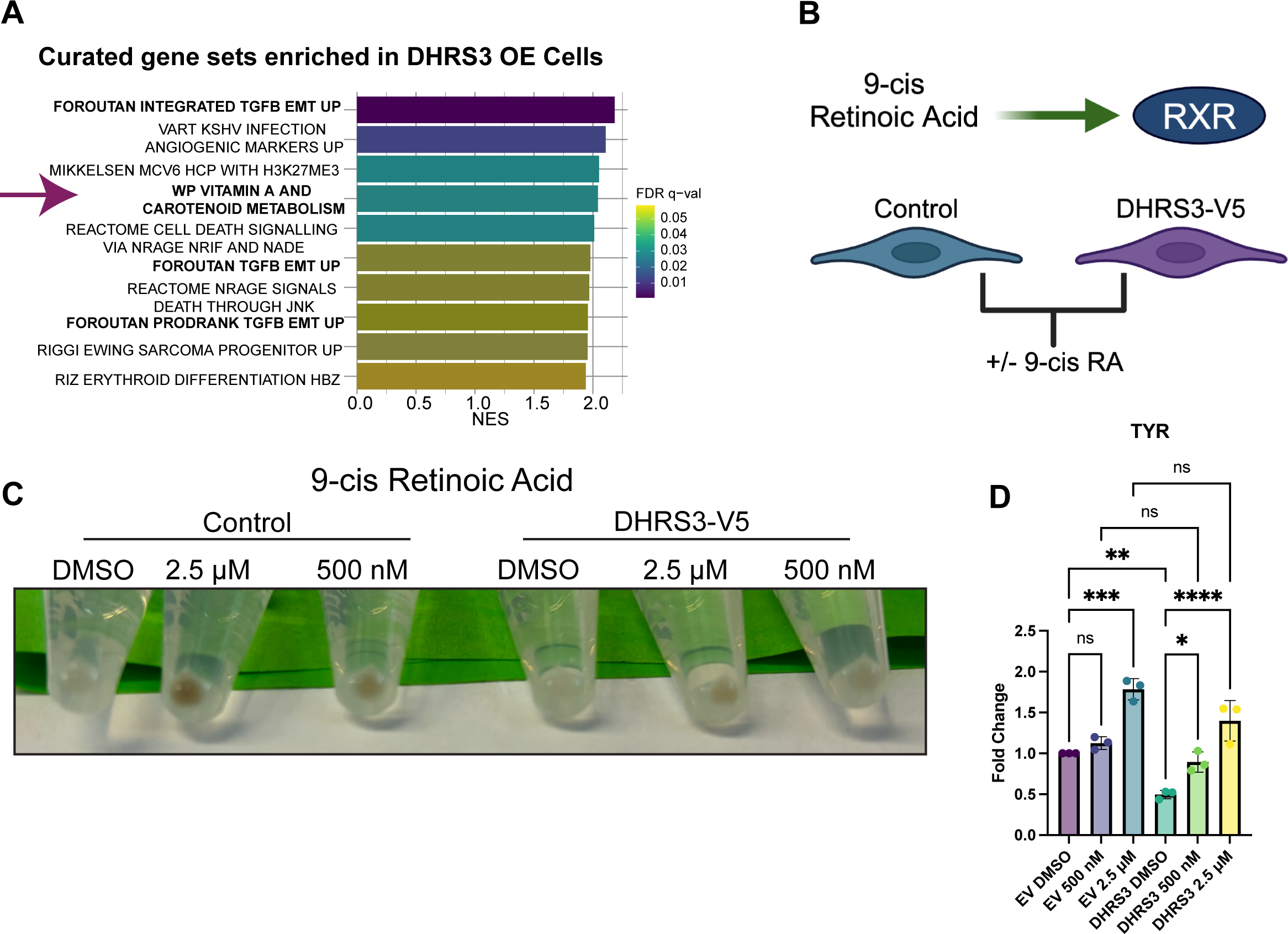
DHRS3 regulates differentiation through retinoid signaling. A. GSEA results using normalized counts from bulk RNA-seq and the Curated Gene Sets list from MSigDB. Retinoic acid related pathway is highlighted by the arrow. B. Schematic of the 9-cis-RA experiment that is shown in C. C. Cells were treated with indicated doses of 9-cis-RA for 3 days and then collected. 250,000 cells per condition were collected, washed in PBS, and imaged to assess pellet color. Image is representative of n = 3 independent experiments. D-G. qPCR of TYR from cells treated as in A. Data were collected in triplicate for n = 3 independent experiments. Graphs are shown with mean +/− SEM and statistics via One-way ANOVA with Holm-Sidak’s multiple comparison correction. * p < 0.05, ** p < 0.01, *** p < 0.001, **** p < 0.0001. Diagrams made with Biorender.com.

DHRS3 functions in an intermediate step of retinoic acid metabolism. It acts in opposition to RDH10, converting all-trans-retinal (which is converted to active retinoic acid) to all-trans-retinol, which can be esterified and stored in lipid droplets (Figure 3B).^25^ Retinoic acid metabolism has long been known to influence melanocyte differentiation and retinoids have been explored as treatment options in melanoma.^27,28,36^ We hypothesized that higher DHRS3 levels would decrease the level of retinoic acid in the cell. This decrease would in turn downregulate RAR/RXR transcriptional activity and in part explain the change in cell state we observe in DHRS3 overexpressing IGR37 cells. To explore the link between DHRS3 levels, retinoic acid metabolism, and cell state we treated IGR37 cells with 9-cis-Retinoic Acid (9-cis-RA), a well-known agonist of retinoid X receptors (Figure 6B).^37^ In line with previous data,^38^ 9-cis-RA treatment causes a significant increase in pigmentation in control cells (Figure 6C). However, consistent with our hypothesis, DHRS3^OE^ cells treated with an equivalent dose of 9-cis-RA were significantly less pigmented than control cells (Figure 6C). This suggests that DHRS3 is acting antagonistically to retinoic acid mediated activation of RXRs. Echoing the effect on pigmentation, when we performed qPCR analysis of Tyrosinase (*TYR*), a key enzyme required for melanin synthesis and pigmentation, we observed a significant decrease in *TYR* expression in DHRS3^OE^ cells. Treatment with 9-cis-RA could rescue this defect, but *TYR* expression remained somewhat lower than in control cells also treated with 9-cis-RA (Figure 6D). This would suggest that DHRS3 antagonizes the effects of RXR activation, potentially by shuttling retinoic acid intermediates towards retinyl ester formation and sequestration in the lipid droplet (Figure 7H).

**Figure 7.**
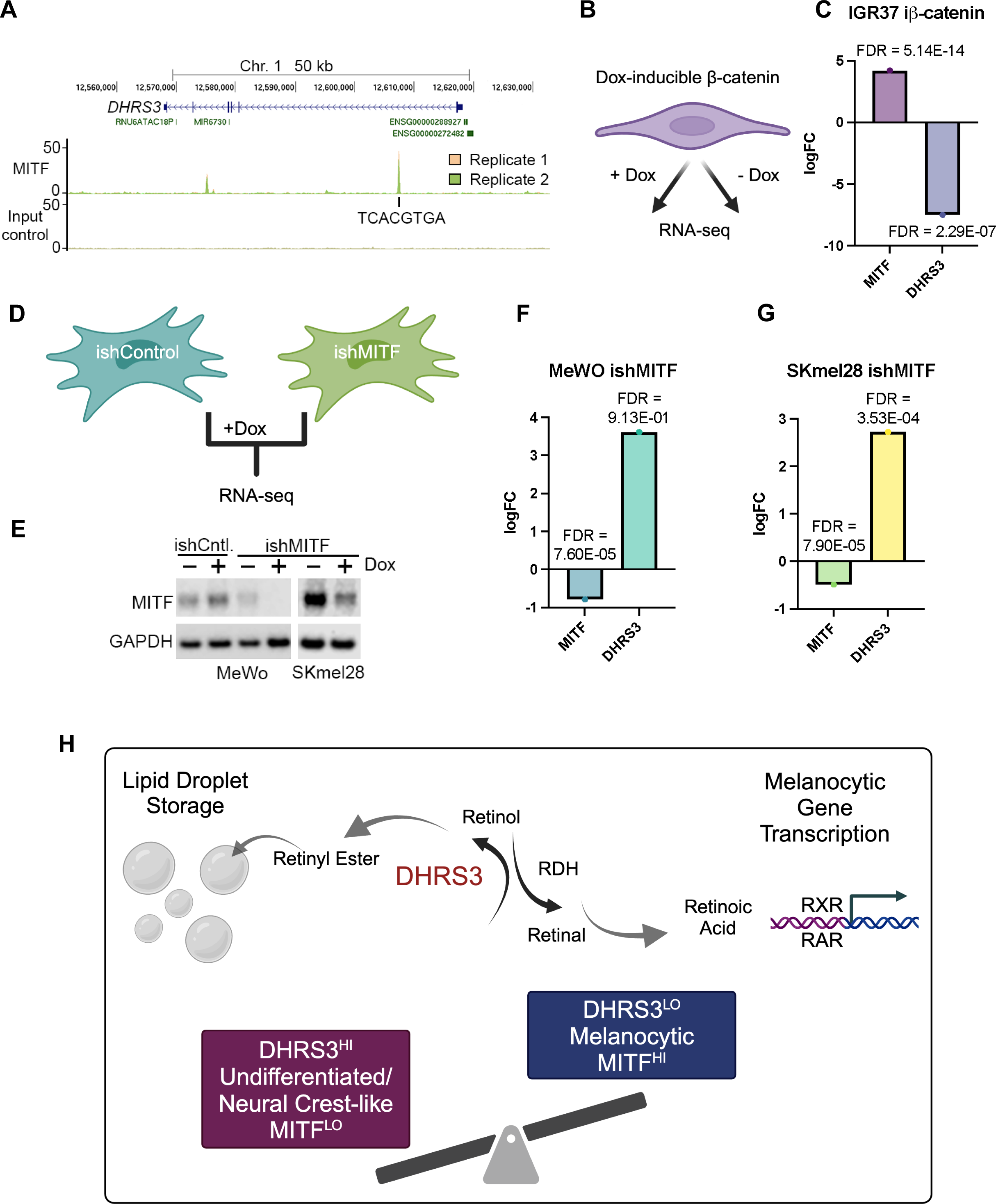
DHRS3 is regulated by MITF, the key regulator of melanoma cell plasticity. A. ChIP-seq data for MITF. The DHRS3 locus is shown and the 8 bp binding site of MITF to Exon 1 of DHRS3 is highlighted.^41^ B-C. Schematic and results of RNA-seq from iβ-catenin cell lines. RNA-seq performed in triplicate. Graph shows the fold change as calculated with EdgeR (see methods). D. Schematic of set up for RNA-seq of MeWO and SKmel28 MITF KD. E. Western blot validating MITF expression levels in the indicated cell lines and conditions. F-G. Graphs showing RNA-seq results for MITF and DHRS3 in the indicated cell lines and conditions. H. Model for DHRS3 role in melanoma cell state. High levels of DHRS3 promote conversion of retinal to retinol. This increases the shuttling of retinoid intermediates away from retinoic acid formation and towards storage as retinyl esters in the droplet. As a consequence, RAR/RXR transcriptional activity is decreased, and cells acquire a more undifferentiated cell state. Diagrams made with Biorender.com.

### DHRS3 is regulated by MITF, the key regulator of melanoma cell plasticity

It has long been observed that melanoma cell state is regulated by MITF, a transcription factor that acts as the master regulator of melanocyte development.^39^ In normal melanocyte development, undifferentiated/neural crest cells express low levels of *MITF* and are unpigmented. As these cells differentiate, MITF activity increases and directly activates expression of pigmentation genes such as Tyrosinase (*TYR*) and *PMEL*.^40^ In melanoma, a similar cascade is maintained, with the *MITF^LO^* cells representing the undifferentiated/neural crest-like state, and the *MITF^HI^*cells representing the melanocytic state.^1^ Given that we found high levels of DHRS3 mRNA and protein only in the undifferentiated/neural crest-like cells, this raised the question of whether DHRS3 expression is regulated by MITF.

To test this, we interrogated a previously published MITF ChIP-seq data set.^41^ This analysis revealed an 8 bp MITF consensus sequence^42^ within the first DHRS3 exon (Figure 7A). As DNA binding is not necessarily indicative of regulation, we next determined whether DHRS3 could be regulated by β-catenin, a known regulator of *MITF* expression^43–45^ and a MITF co-factor.^46^ We used IGR37 human melanoma cells engineered to express doxycycline-inducible β-catenin^47^ followed by RNA-seq of cells treated with doxycycline compared to untreated cells (Figure 7B).^47^ In IGR37 cells overexpressing β-catenin, we quantified a marked 18-fold up(+4.2log2) up-regulation of *MITF* expression and a dramatic reduction (128-fold, −7log2) in *DHRS3* mRNA expression (Figure 7C, Supplementary Table 6). This result suggests that DHRS3 is repressed either by β-catenin complexed with other DNA binding factors such as LEF/TCF, or by MITF itself. To test for regulation by MITF, we used two human melanoma cell lines, MeWo and SKmel28, engineered to express doxycycline inducible shRNA for MITF (Figure 7D-E). Eight days after induction of MITF-targeting shRNA or a control non-targeting, we performed bulk RNA-seq (Figure 7D, F). The results were consistent with MITF repressing *DHRS3* expression, with a 12-fold (+3.2 log) increase in *DHRS3* expression in MITF depleted MeWo cells, and a 6.6-fold (+2.72 log2) increase in *DHRS3* expression in MITF depleted SKmel28 cells (Figure 7F-G, Supplementary Table 6). Together with the ChIP-seq result showing binding of MITF with the first intron of *DHRS3*, these data strongly suggest that DHRS3 is directly repressed by MITF. This is consistent with previous observations^48^ that revealed repression of genes by MITF is rarely via promoter binding. Collectively, this data demonstrates that expression of DHRS3 is directly controlled by MITF, and that in turn high levels of DHRS3 support the undifferentiated/neural crest-like cell state by affecting expression of genes linked to de-differentiation and retinoic acid metabolism.

## Discussion

It is now well recognized that tumors exist in a large number of distinct transcriptional states.^49^ While such heterogeneity has been observed for decades, the advent of single-cell RNA seq has provided further nuance to defining these states.^6,50^ In melanoma, these different transcriptional states have important practical consequences in that each state corresponds to a distinct cellular phenotype. For example, cells with a low level of MITF are more invasive and drug resistant, whereas cells with high levels of MITF are more pigmented but also more proliferative.^1–5,7,51^ Dynamic switching between these states is known to occur, but both the upstream triggers as well as the downstream effectors of switching remain intense areas of investigation.

Lipids are increasingly being investigated as mediators of cell state plasticity in cancer. Because lipids are strikingly diverse in structure and function, each species can act at multiple levels to mediate changes in cellular physiology. Increased fatty acid uptake, mediated by molecules such as CD36 or the SLC27A/FATP family of proteins, can render the cancer cell more invasive and drug resistant across multiple tumor types.^14,52–55^ These fatty acids can be used for diverse processes including incorporation into membranes, β-oxidation, or conversion into acetyl-CoA and subsequent protein acetylation.^12,15,56–60^ Lipids can also play a key role in ferroptosis, a central mediator of cancer cell survival. Dependent on fatty acid saturation, excess lipids can promote ferroptosis through lipid peroxidation while uptake of monounsaturated fatty acids can protect cancer cells from ferroptosis.^8,61–63^

Less well studied is the role of the lipid droplet itself. In both de novo lipid synthesis as well as after uptake from extrinsic sources, lipids such as fatty acids are stored in these specialized organelles. But this organelle is not simply an inert storage container. The droplet itself plays a role in the physiology of both normal and cancer cells.^64,65^ In part, this is because the lipid droplet makes close contact with many other organelles within the cell, including the ER, mitochondria, peroxisomes, and lysosomes.^66^ The specific role of lipid droplets in cancer is less well understood, but they likely play diverse roles in processes such as preventing lipotoxicity, buffering nutrient deprivation, metabolic flux, signaling, and stemness.^67–75^

While many of the above studies have focused on the lipid content within the droplet, an increasing number of studies have begun to characterize the diverse array of proteins found on the surface of the lipid droplet membrane. In the past few years, there have been meaningful advances in generating high confidence lipid droplet proteomes. For example, a recently developed proximity labeling approach in U2OS osteosarcoma and Huh7 hepatocellular carcinoma cells generated a high confidence list of LD proteins^20^ revealing both expected but also unexpected components of the membrane. At a functional level, genetic perturbation of lipid droplet proteins such as DGAT1^12,13,76,77^ or PLIN2^78–80^ can affect many canonical hallmarks of cancer such as proliferation, invasion, metastasis and drug resistance.

A major unanswered question is whether these lipid droplet proteins are simply a reflection of cell state, or whether they can act as drivers of cell state. In our study, we find that DHRS3 itself can enact a change in melanoma cells towards a more undifferentiated state. However, this effect is tightly linked to the way that DHRS3 interacts with the lipid content of the droplet. DHRS3 and RDH10 act in opposition to each other,^25^ and overall regulate the relative levels of retinal versus retinol. In the presence of high levels of DHRS3, there is less active retinoic acid in the cell, which therefore makes the cell less differentiated and pigmented. Rescuing this deficiency via exogenous retinoic acid restores pigmentation and promotes differentiation. Our data are consistent with a model in which DHRS3 levels are linked to melanoma cell state through the balance of retinoic acid within the lipid droplet (Figure 7H). However, one unresolved question in our work is the observation that drug resistant, neural crest-like cells have previously been shown to have high levels of RXRG signaling, and that treatment with RXR antagonists can delay such resistance.^2^ It is probable that RXR signaling can play different roles in homeostatic conditions (as in our study) versus after the pressure of drug therapy. We do not fully understand how retinoic acid levels would differ in these unique conditions, but one possibility is that the binding partners of RXR could differ. Delineating these discrepancies should be a major focus of future studies.

A key finding of our study is the observation that DHRS3 is repressed by MITF, the master regulator of melanoma cell plasticity. While it is clear that MITF can shift the balance between the undifferentiated vs. differentiated state, what is less clear is the way in which it regulates other aspects of cell state. Aside from differentiation and pigmentation MITF also regulates cell cycle progression through an interaction with Rb and p21.^81^ Our data suggests a new role for MITF in cellular plasticity, which is repression of DHRS3, which in turn regulates lipid droplet composition. It further demonstrates the complexity of the ways in which MITF can act as both a transcriptional activator and repressor, which in part is determined by its interaction with β-catenin.^46^

While our study initially characterized the lipid droplet proteome in A375 cells, a limitation of our study is that there could be significant variation in the lipid droplet proteome in distinct melanoma cell states. DHRS3 is one such example, since it is highly enriched in the undifferentiated/neural crest-like cell state, and not very well expressed in more differentiated cells. Given this diversity, future studies should aim to characterize the proteome more broadly across a range of cell states. In addition, our study is limited by the fact that we were only able to characterize the proteome in cell lines, as opposed to *in vivo* tumors. Repeating these studies *in vivo* would be a major challenge, as the number of cells required to isolate the droplets and then perform mass spectrometry is large. However, with the advent of increasingly sensitive mass spectrometry methods, this may be overcome in the near future. Finally, an unanswered question in our study is the function of the other lipid droplet proteins (aside from DHRS3) that we have identified. Some of these proteins are shared with other compendium lipid droplet proteomes from different cell types, but others appear to be somewhat unique to melanoma. Future studies should aim to systematically investigate the function of these other proteins in different melanoma cell contexts to understand whether they directly affect melanoma cell state or play other roles in the disease.

## Acknowledgements

We thank the Rockefeller Proteomics Resource Center for their technical support for LC-MS/MS experiments. The Molecular Cytology Core Facility provided confocal imaging support. We also thank all members of the White lab for helpful discussions. EJ was supported by a F31 Ruth L. Kirschstein Predoctoral Individual National Research Service Award (F31AR079215) from the National Institute of Arthritis and Musculoskeletal and Skin Diseases. MVH was funded by a K99/R00 Pathway to Independence Award from the National Cancer Institute (K99CA266931). CRG was funded by the Ludwig Institute for Cancer Research and CRG and PL by NIH R01 CA268597-01. YM was supported by a Medical Scientist Training Program grant from the NIH under award number T32GM007739 to the Weill Cornell/Rockefeller/Sloan Kettering Tri-Institutional MD-PhD Program and Kirschstein-National Research Service Award (NRSA) predoctoral fellowship under award number F30CA265124. R.M.W. was funded through the NIH/NCI Cancer Center Support Grant P30 CA008748, the Melanoma Research Alliance, The Debra and Leon Black Family Foundation, NIH Research Program Grants R01CA229215 and R01CA238317, NIH Director’s New Innovator Award DP2CA186572, The Pershing Square Sohn Foundation, The Mark Foundation for Cancer Research, The American Cancer Society, The Alan and Sandra Gerry Metastasis Research Initiative at the Memorial Sloan Kettering Cancer Center, The Harry J. Lloyd Foundation, Consano and the Starr Cancer Consortium (all to R.M.W.).

## Author Contributions

EJ and RMW developed the experiments and interpreted the results. EJ performed all experiments and analysis unless otherwise noted. YM performed RNA-sequencing analysis of IGR37 DHRS3 cells. MVH and JR assisted with computational analyses. CP and HM helped with experimental design and execution of LC-MS/MS and performed the respective data analysis. PL and CRG performed ChIP-seq analysis and RNA-seq experiments of iβ-Catenin and ishMITF cells. EJ and RMW wrote the manuscript. RMW acquired funding for the project. All authors read and edited the manuscript.

## Star Methods

### Lead Contact

Richard M. White

### Data and Code Availability

Raw and processed RNA-seq and Proteomics data generated in this study have been deposited in Gene Expression Omnibus (GEO) under GSE262095 and PRIDE under PXD050693. Previously generated RNA-seq of iβ-Catenin and ishMITF cells are available under accession code GSE136803 and GSE255256 respectively. Re-analyzed scRNAseq data are available under GSE134432. ChIP-seq data is available on GEO under GSE132624. Code used in this study has been deposited on Github at github.org/emj464/Johns-2024

### Materials Availability

All other relevant data and materials supporting key findings in this study are available within the article or from corresponding authors upon reasonable request.

## Key Resources Table

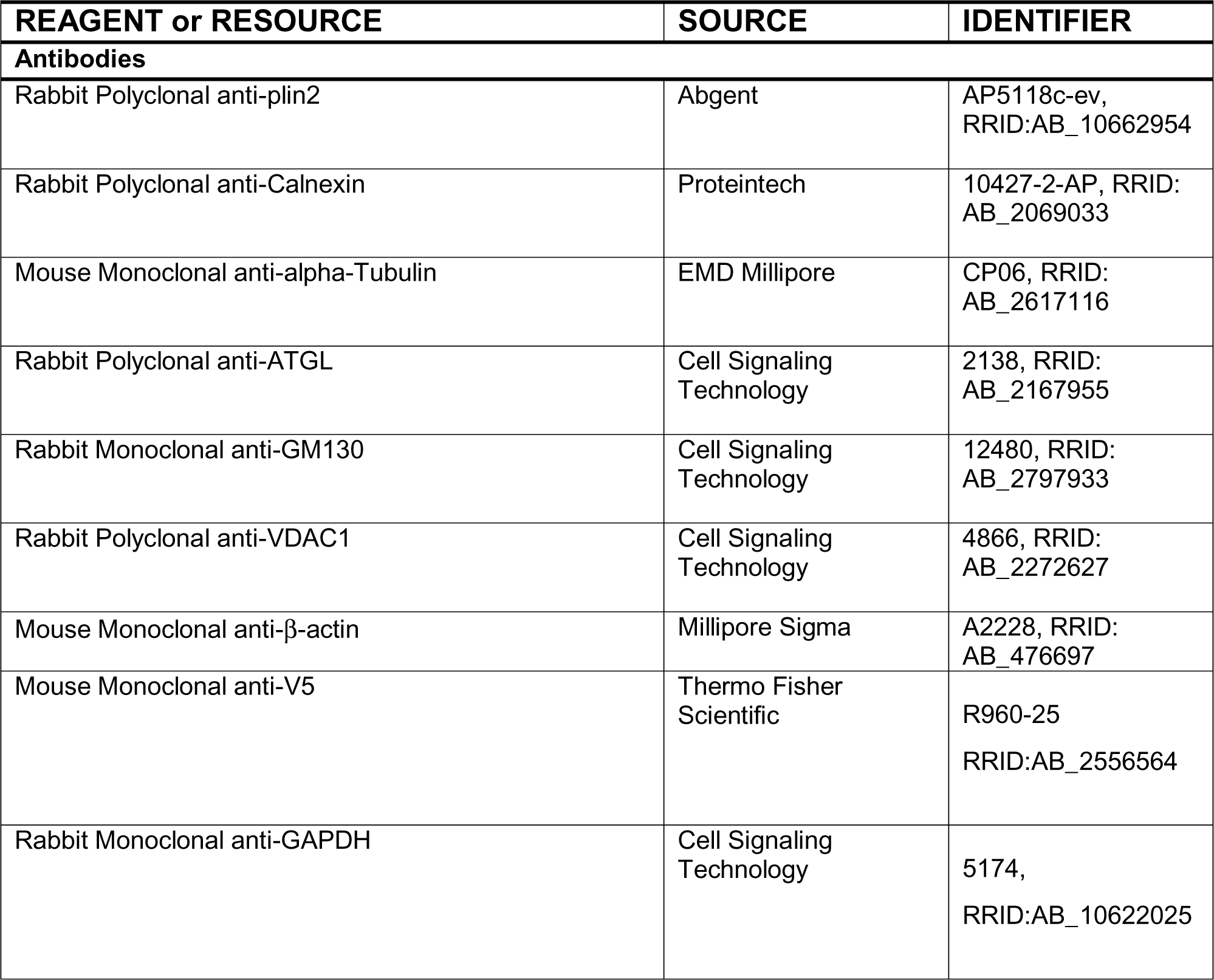
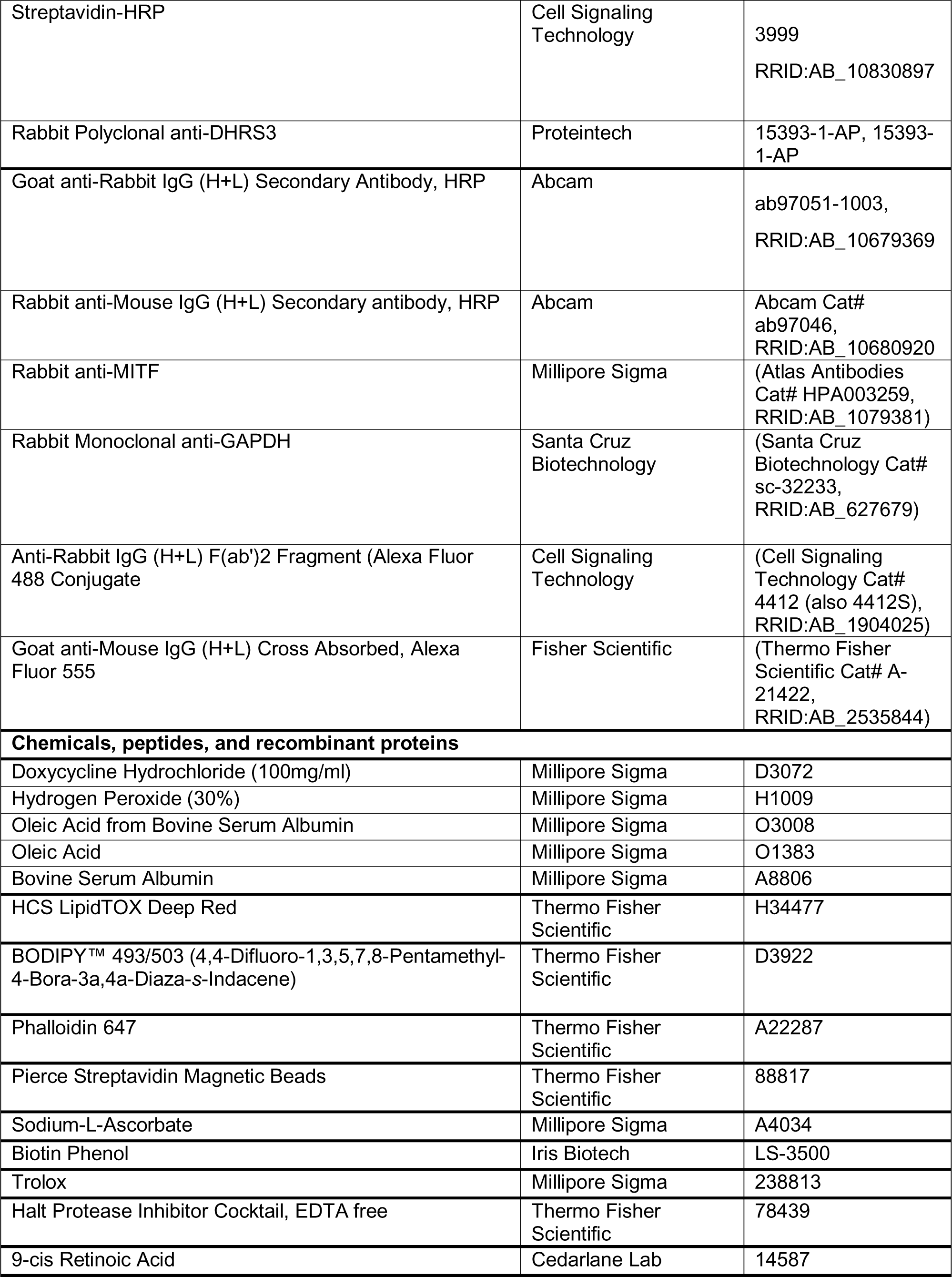
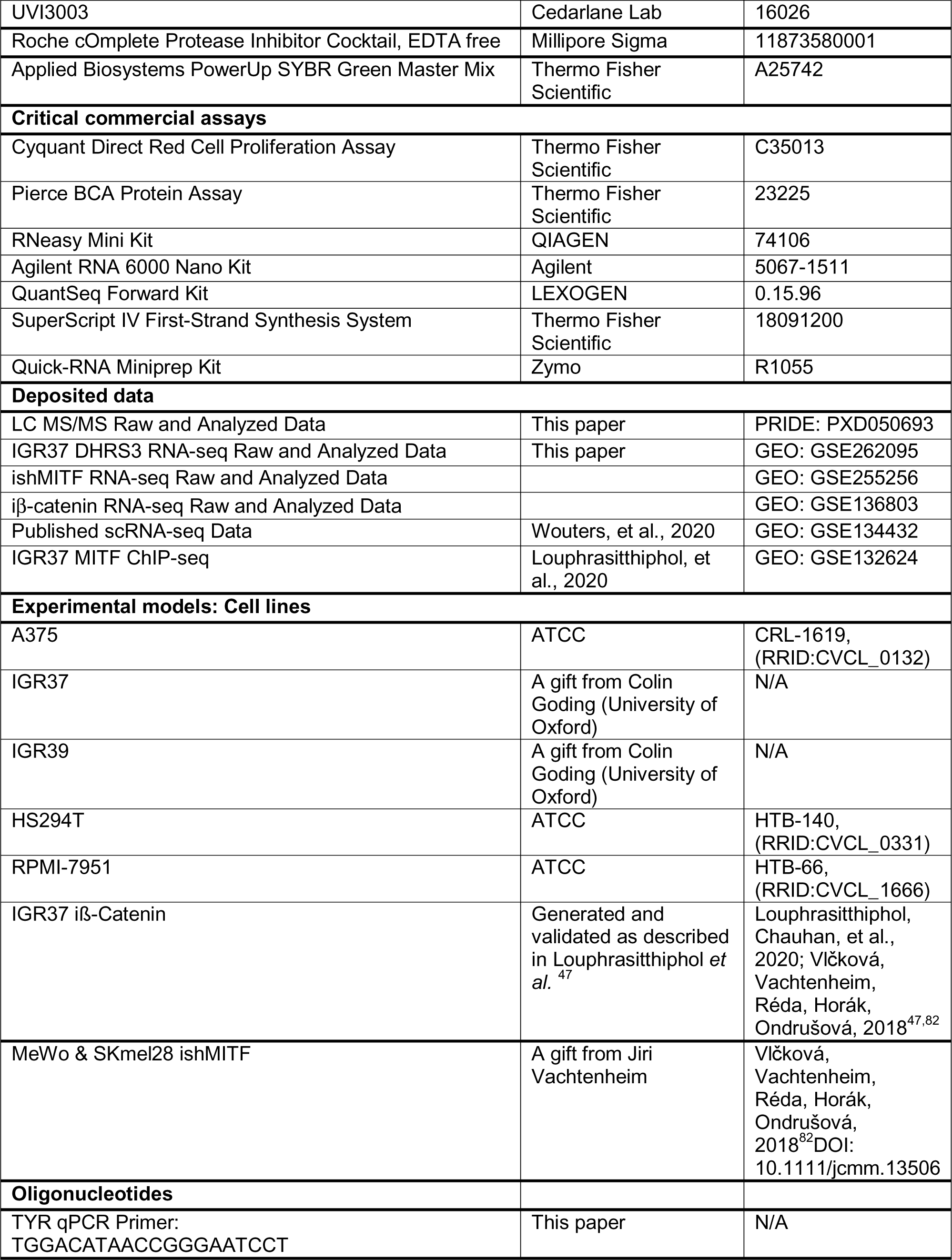
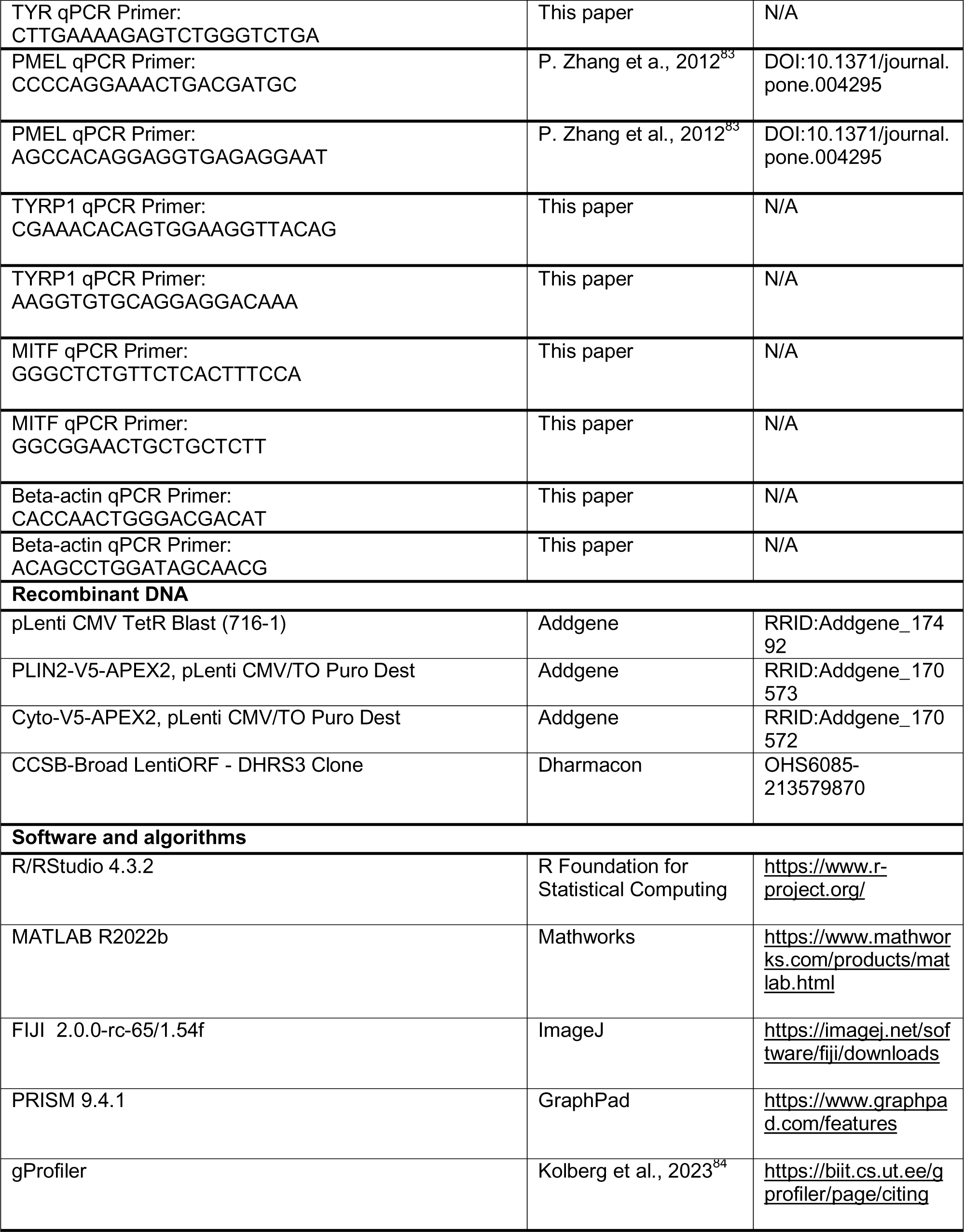
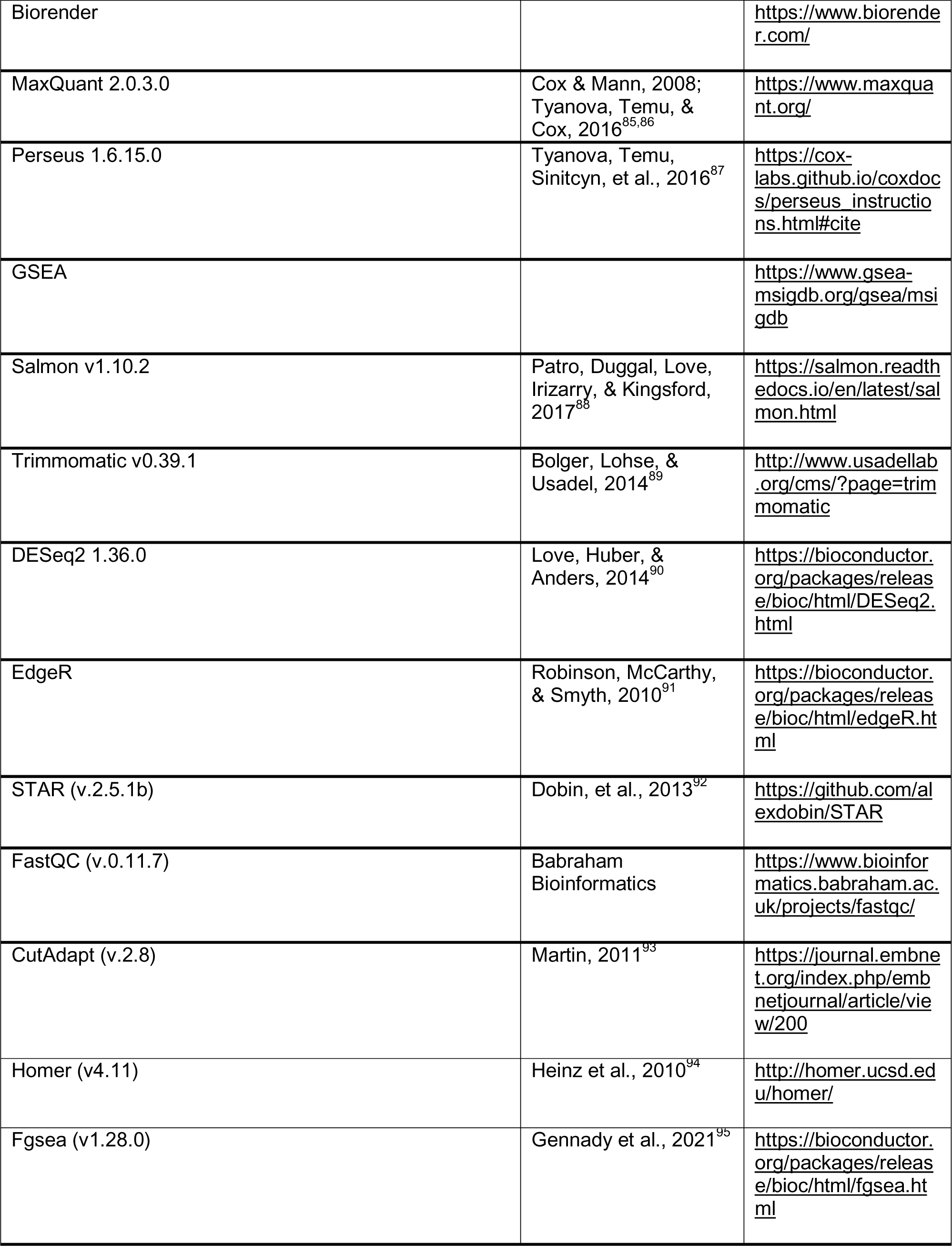

### Cell Culture

A375, HS294T, and RPMI-7951 cells were obtained from ATCC and routinely tested for Mycoplasma. IGR37 and IGR39 cells were a gift from the Colin Goding lab. Upon receipt cell line identity was confirmed by STR profiling and cells were tested for Mycoplasma (09/2023). All cells were maintained in a 37°C and 5% CO_2_ humidified incubator. A375, HS294T, and RPMI-7951 cells were maintained in DMEM (Gibco, 11965) supplemented with 10% FBS (Gemini Bio, 100-500). IGR37 and IGR39 cells were maintained in RPMI (Gibco, 11875119) supplemented with 10% FBS. Where indicated cells were treated with pre-conjugated Oleic Acid (Sigma Aldrich #O3008) or Oleic Acid:BSA as described below.

APEX2 expressing A375 cells were generated as follows. A375 cells were infected with pLenti CMV-TetR Blast virus (Addgene #17492) with 4 µg/ml polybrene followed by selection with 10 µg/ml Blasticidin. A375 TetR expressing cells were subsequently infected with PLIN2-v5-APEX2 (Addgene #170573) or Cyto-v5-APEX2 (Addgene #170572) with 4 µg/ml polybrene followed by selection with 1 µg/ml Puromycin.

IGR37 DHRS3 overexpression cells were generated as follows. IGR37 cells were infected with either pLX304 empty vector virus (Addgene #25890) or CCSB-Broad LentiORF-DHRS3 Clone (Clone ID: ccsbBroad304_07379) virus with 4 µg/ml polybrene followed by selection with 10 µg/ml Blasticidin.

MeWo Cells were maintained in EMEM (ATCC) with 10% FBS while SKmel28 cells were maintained in RPMI-1640 (GIBCO) with 10% FBS (Biosera) Both were grown in a humidified incubator at 37°C and 10% CO_2_.

### Western Blotting

For APEX2 expressing cells, cells were lysed in HLM buffer (20mM Tris-HCl, pH 7.4 and 1 mM EDTA, 1x Halt protease inhibitor cocktail) supplemented with 1% SDS and sonicated at 10% power with 10 1 second pulses. Following sonication, lysates were boiled for 5 min at 65°C and spun down for 10 mins at 13,000 RPM at RT. Protein concentration was measured with the Pierce BCA Assay and equal amounts of protein (in µg) were mixed with 1X Laemmli Sample Buffer (#BP-111R) and boiled at 95°C for 5 mins. For DHRS3 westerns, cells were lysed in 1% SDS (50mM Tris-HCl, 150mM NaCl, 2.5mM EDTA, 1x Halt protease inhibitor cocktail). Lysates were boiled for 5 minutes and then spun down for 10 mins at 13,000 RPM at RT. Protein concentration was measured as above.

Equal amounts of protein (10-40 ug) were loaded onto 4-15% Mini-PROTEAN TGX Precast Protein Gels for electrophoresis at 120 volts followed by semi-dry electrophoretic transfer to nitrocellulose membranes via the Bio-Rad Trans-Blot Turbo transfer system. Membranes were blocked in 5% nonfat milk/TBS-T (0.1% Tween-20) at RT for 1 hour. Membranes were then incubated in primary antibody overnight at 4 °C in either 5% BSA or 5% Milk in TBST followed by 3x 5 min washes at RT. Secondary antibodies were added (1:10,000) in 5% milk in TBST for incubation at RT for 1 hour follow by 3x 5 min washes. ECL substrate (Cat #32106 or #WBKLS0500). Visualization was carried out on an Amersham Imager 600. Antibody dilutions: PLIN2 (Abgent AP5118c-ev) 1:1000, Calnexin (Proteintech 10427-2-AP) 1:20,000, Tubulin (EMD Millipore CP06) 1:1000, ATGL (CST 2138) 1:1000, GM130 (CST 12480) 1:1000, GAPDH (CST 5174) 1:5000, VDAC1 (CST 4866) 1:1000, Actin (Sigma A2228) 1:10,000, V5 (Thermo Fisher R960-25) #1:5000, Streptavidin HRP (CST 3999) 1:2000, DHRS3 (Proteintech 15294-1-AP) 1:500.

For Figure 5H Treated cells were lysed directly in 4x LDS (Invitrogen, # NP0008) supplemented with 10% v/v β-mercaptoethanol, heated at 95 °C for 10 min before SDS-PAGE using Bis-Tris gels (Invitrogen, #WG1403) Gels were transferred onto PVDF Membrane (Amersham) at 100 V for 60 min for Western-blotting, blocked 1 h, room temperature in 5% non-fat milk in TBS containing 0.25% TWEEN-20 (TBST). Primary antibody incubations were done in 5% BSA-TBST overnight and 1 h RT for secondary antibodies. Visualization was carried out on X-ray film (Fujifilm) using ECL (Amersham). Antibodies used were MITF (RRID:AB_1079381) and GAPDH (RRID: AB_627679).

### Immunofluorescence Staining

Cells were seeded in either a Millicell EZ 4-well chamber slide (#PEZGS0416) or #1.5 coverslips (EMS #50-192-9522) coated with Poly-L-lysine (Millipore Sigma #P4707). After washing with PBS, cells were fixed for 15 minutes in 4% Formaldehyde (Fisher Scientific #FB002) at room temp. Cells were permeabilized with either 1% Triton or 0.01% Digitonin (for DHRS3) for 30 min at RT. For antibody staining, cells were washed 3x in PBS and then blocked in 10% normal goat serum (Thermo Fisher #50-062Z) for 1 hr at RT. Primary antibodies were added at the indicated concentrations in 10% goat serum and incubated overnight at 4°C. Samples were washed 3x in PBS and then incubated with the appropriate secondary antibodies at 1:500 in 10% goat serum for 1 hr at RT. If using, BODIPY 493/503 (Thermo Fisher D3922) was added to the secondary antibody solution at 1:100. Where indicated, LipidTOX Deep Red 1:1000 (Invitrogen #H34477) in PBS was added to samples to stain LDs for 30 mins at RT. After staining with secondaries/LD dyes, samples were washed 3x in PBS and incubated with Hoechst 1:2000 (Thermo Fisher #H3570) for 5 mins at RT. Samples were then washed an additional 3x times in PBS before mounting on slides in Vectashield (Vector Laboratories #H-1900-10). If no antibody staining was required, cells were permeabilized, washed 3x in PBS and then incubated with PBS containing 1% BSA, BODIPY 493/503 1:100, and Phalloidin 647 1:200 (Thermo Fisher A22287) for 1 hour at RT. This was followed by staining with Hoechst and mounting as above. Antibody concentrations: V5 (Thermo Fisher R960-25) 1:500, Streptavidin-594 (Thermo Fisher S32356) 1:500, DHRS3 (Proteintech 15294-1-AP) 1:200, Tubulin (EMD Millipore CP06) 1:1000. Samples were imaged with the Zeiss LSM880 inverted microscope with 63x oil immersion objective. Where needed, Airyscan was used to increase resolution. Confocal z-stacks were visualized in Fiji.

### APEX2 Labeling & Lipid Droplet Isolation

LD labeling & isolation was performed as previously described with some modifications.^20,24^ For LD isolation 4 million cells were seeded in each 245 mm square plate with 4 plates per condition. 24 hours after seeding, cells were treated with Dox at 5 ng/ml. 24 hours before collecting, Oleic Acid (200 μM) + Dox 5 ng/ml was added to the media. For APEX2 labeling, 500 μM Biotin Phenol was added to cells and incubated at 37°C for 30 mins. Then, 1 mM H_2_O_2_ was added to the media and incubated for 1 min at RT. The reaction was quenched with 3x washes in Quenching Buffer (5mM Trolox, 10mM Sodium Ascorbate in DPBS) followed by a final wash in PBS. Cells were scraped in PBS on ice and transferred to a 50 ml conical tube followed by a spin at 500g, 4°C for 10 mins. Cells were resuspended in 3ml HLM buffer (20mM Tris-HCl, pH 7.4 and 1 mM EDTA, 1x Roche cOmplete Protease inhibitor cocktail), incubated for 10 mins on ice, and then transferred to a Dounce Homogenizer. Cells were homogenized with 80x strokes of the tight fitting Dounce Homogenizer, followed by a spin at 1000xg for 10 mins at 4°C. Between 2-3 ml of Post Nuclear Supernatant (PNS) was transferred to a Beckman Open-Top Thinwall Ultra-Clear Tube (#344059). The PNS was diluted to a final concentration of 20% sucrose, then 5 ml of 5% sucrose was layered on top, followed by 3.5-4.5 ml HLM (0% sucrose). Tubes were spun at 28,000g for 30 mins at 4°C in a TH-641 swinging bucket rotor. After centrifugation, lipid droplet fraction (∼1 mL) was sliced off the top of the tube using a tube slicer. The remaining volume was collected in 1 mL fractions to assess the quality of the fractionation.

To analyze fractionation by western blot, equal percent by volume of each fraction was mixed with 10% SDS to a final concentration of 1% SDS. For the lipid droplet fraction, samples were incubated at 37°C and sonicated 3x 10 seconds at 10% amplitude every 20 minutes for a total of 1 hour. This was followed by a final incubation for 10 mins at 65°C, a spin for 10 mins at 14,000 RPM. The soluble layer was removed and transferred to a new tube. All other fractions were sonicated 3x 10 seconds at 10% amplitude followed by a final incubation for 10 mins at 65°C.

For streptavidin IP and mass spectrometry: Lipid droplet fractions were collected sequentially and stored at −80°C until use. LD proteins were precipitated with 6x volume ice cold acetone at −20°C overnight. Samples were spun down at 14,000 x g at 4°C for 10 mins. Acetone was removed and pellet was allowed to dry (under foil) for 5-10 mins. Pellet was resuspended in RIPA + protease inhibitors + 2% SDS and boiled at 95°C for 10 mins and then spun down to ensure no visible pellet for 10 mins at 14,000 RPM. Equal amounts of protein per sample were then diluted in SDS free RIPA to a final 0.1% SDS concentration and then incubated with 25 ul solubilized, samples were diluted in SDS free RIPA buffer (0.1% final SDS) and then incubated with Pierce Streptavidin Magnetic Beads (cat #88817) overnight at 4°C in an end over end rotator. The next day, beads were washed for 2 min at RT (unless otherwise indicated) with the following: 2x RIPA buffer (SDS free) buffer, 2x NP-40 buffer(1% NP-40), 1x 1M KCl, 1x 0.1 M Sodium Carbonate (10 seconds), 1x 2 M Urea, (10 seconds), 3x 10mM Tris-HCl, pH8.0 (changing tubes each time). At the final wash 100 ul of buffer/beads was removed (10% of sample) for WB analysis of IP while the rest was used for mass spectrometry.

### Lipid Droplet Proteomics by Mass Spectrometry

Dried magnetic streptavidin beads were suspended in a solution 25uL of 50 mM Ammonium Bicarbonate/10 mM DTT for reduction of proteins and incubated for one hour at room temperature. Reduction was followed by alkylation with the addition 7.5uL of 100mM Iodoacetamide/50 mM Ammonium Bicarbonate and incubated for one hour at room temperature in the dark. Beads then underwent partial on-bead digestion with 250 ng of Porcine Trypsin (Promega) in 50 mM Ammonium Bicarbonate for three hours at room temperature. Supernatant was then extracted to undergo digestion with 500 ng of Porcine Trypsin (Promega) and 500 ng of Endopeptidase Lys-C (Wako) in 50 mM Ammonium Bicarbonate overnight at room temperature. Digestion was halted by addition of neat trifluoroacetic acid. Peptides underwent reversed phase based micro solid phase extraction.^96^ 3 of 10uL were injected and analyzed by nano LC-MS/MS (Fusion LUMOS coupled to an Easy-nLC 1200, Thermo Scientific). Mass spectrometer were mass calibrated weekly and operated with lock mass.^97^ MS and MS/MS were recorded at a resolution (@ 200 Th) of 120,000 and 30,000, respectively. Automatic Gain Control (AGC) was set to 4e5 and 5e4 for MS and MS/MS. Samples were analyzed using a gradient increasing from 2% B/98% A to 35% B/65% A in 70 min (A: 0.1% formic acid, B: 80% Acetonitrile/0.1% formic acid). Peptides were separated using a 12cm/100um packed in column emitter (NIKKYO TECHNOS CO., LTD.).

### Analysis of Mass Spectrometry Data

Data were queried against the database: Uniprot_human_June_2022 concatenated with common contaminants.^98^ using MaxQuant^85,86^ v 2.0.3.0. In short: 20 ppm mass accuracy was used for precursor mass accuracy and 20 mDa for fragment ions. Carbamidomethylation of cysteines was set as a fixed modification and oxidation of methionine and protein N-termini acetylation were set as variable modifications. Search results were filtered using false discovery rates of 2% or better for peptides and 1% or better for proteins. Intensity-Based Absolute Quantitation (iBAQ) values^99^ were used to relative quantify matched proteins. Utilizing Perseus^87^ software platform (v 1.6.15.0), iBAQ values were log2 transformed, filtered for common contaminants and proteins found via reverse search. The data was further filtered by requiring signals in at least 75% of replicates for at least one of the conditions, resulting in 407 proteins being quantitated. Missing values were imputed. In addition, to generate normalized values, the average (non-log2 transformed) Cyto-APEX2 iBAQ value for each protein was subtracted from the average PLIN2-APEX2 iBAQ values as previously described.^24^ Proteins were then ranked from highest to lowest normalized iBAQ value. To generate a high confidence list of lipid droplet proteins, we set a manual cutoff such that approximately 60% of previously validated lipid droplet proteins were identified. This yielded a list of 96 LD proteins total. To generate the heatmap, Log_2_ normalized iBAQ values with imputation are shown across all 10 conditions.

To test for enrichment of lipid droplet proteins, we ran g:Profiler^84^ with GO:Cellular Components and Biological Processes using the high confidence lipid droplet protein list. This revealed significant enrichment of lipid droplet and related pathways.

### Oleic Acid:BSA Preparation

For proteomics experiments, Oleic Acid:BSA was prepared as follows. Stock solutions of Oleic Acid (Sigma #O1383) were resuspended in ethanol and aliquoted. To complex; 9 mM Oleic Acid was mixed with 3 mM BSA (Sigma #A8806) in PBS (3:1 ratio) and incubated at 37°C in a shaker for 2 hours until the solution cleared. OA:BSA was then sterile filtered, aliquoted, and stored at −20°C until use.

### Re-analysis of publicly available melanoma -omics data

CCLE and TCGA RNA-seq data was downloaded and analyzed as previously described.^12^ CCLE Proteomics data was downloaded from https://depmap.org/portal/download/custom/ and z-scores were used to compare protein levels across different cell states as above.

Single-cell RNA-seq data from human melanoma patients^5^ was downloaded from GEO (GSE134432). All analysis was done in R (v. 4.3.1) using the Seurat^100^ package (version 5.0.2). The counts matrices for each cell line were imported into R and converted into Seurat objects using the function CreateSeuratObject with default parameters. The Seurat objects were then merged into one combined object containing data for all cell lines. Using the merged object, counts were normalized with SCTransform.^101^ UMAP was calculated using the Seurat function RunUMAP with 15 principal components.

### Bulk RNA-sequencing and analysis

For RNA-seq of IGR37 DHRS3 OE cells, 3 replicates of 1 million cells each were seeded in 10 cm dishes. Cells were allowed to grow for 2 days and then collected for RNA-seq and snap frozen. Library preparation and sequencing was done by Azenta Life Sciences. FASTQ reads were trimmed with Trimmomatic^89^ v.0.39.1, then mapped and quantified with Salmon^88^ v1.10.2 using selective alignment with the GRCh38 reference genome2. Downstream analysis was done in R v4.2 or v4.3.2. Differential gene expression analysis was performed using DESeq2 (v1.36)^90^ with default parameters. Gene set enrichment analysis was done using GSEA^102^ with Hallmark and Curated Gene Set pathways from MSigDB or fgsea.^95^ To run GSEA, we pre-filtered the normalized counts matrix to remove genes with counts <10 and genes with no official gene name. For fgsea, we ran this using the log2FC output from DESeq2 against the custome gene pathways for melanoma cell state.^4^

Known motif analysis was performed with HOMER function findmotifs.pl and searching for motifs 8, 10, and 16 bp in length within +/− 500 bp of the TSS.

For RNA-seq of iβ-Catenin and ishMITF cells: iβ-Catenin cell line was generated, validated and treated as previously described^47^ for total RNA extraction and sequencing as ishMITF cells. ishMITF cell lines and lentiviral vectors were a kind gift from Jiri Vachtenheim^82^. Cells were seeded on 6 cm dish overnight, the next day, fresh media with 1 µg/ml Doxycycline was added. Media with or without Doxycycline was changed every other day. After 8 days, the media was aspirated, and the plates snap frozen at −80 °C before total RNA extraction using RNeasy Mini Kit (QIAGEN #74106) as per manufacturer protocol. RNA quality was assessed on Bioanalyzer 2100 (Agilent) using Agilent RNA 6000 Nano Kit (#5067-1511). Only samples with RIN values of over 9.5 were subjected to library prep and sequencing using the Wellcome Trust genomic service, Oxford. QuantSeq Forward kit (LEXOGEN #0.15.96) was used for library prep, using 500 ng starting material to minimize the PCR amplification step. Samples were sequenced on HiSeq 4000 (Illumina).

Base calling, demultiplexing were carried out by the Wellcome Trust Genomic Service, Oxford. Quality of the raw sequencing fastq files were evaluated using FastQC (v.0.11.7). Adapter contamination removed and poly-A trimmed using CutAdapt ^93^ (v.2.8) before mapping to the human reference genome (hg38) with STAR ^92^ (version 2.5.1b). Gene expression normalization and differential gene expression analyses were performed as previously described^32,33,47^ using edgeR glmQLFTest.

### qRT-PCR

Cells were treated with the indicated RXR agonists and inhibitors for 3 days. After collecting the cell pellet, total RNA was isolated using the Quick-RNA Miniprep Kit (Zymo R1055) according to manufacturer’s instructions. cDNA was synthesized using SuperScript IV First-Strand Synthesis System (Thermo Fisher, 18091200) and qPCR was performed using Applied Biosystems PowerUp SYBR Green Master Mix (Thermo Fisher, A25742). Results were normalized to the *beta-actin* housekeeping gene.

### Statistics and Reproducibility

Statistical analysis and figures were produced using GraphPad Prism, R/RStudio and Biorender.com. Image processing and analysis was performed using Fiji. The statistical analysis and p-values are presented in each respective figure legend. Unless otherwise noted in the legend, data are representative of at least 3 independent experiments.

## Data and Code Availability

Raw and processed RNA-seq and Proteomics data generated in this study have been deposited in Gene Expression Omnibus (GEO) under GSE262095 and PRIDE under PXD050693. Previously generated RNA-seq of iβ-Catenin and ishMITF cells are available under accession code GSE136803 and GSE255256 respectively. Re-analyzed scRNAseq data are available under GSE134432. ChIP-seq data is available on GEO under GSE132624. All other relevant data supporting key findings in this study are available within the article or from the corresponding authors upon request. Original code used in this study will be deposited on Github.

**Excel Table 1: List of previously validated lipid droplet proteins**

**Excel Table 2: iBAQ values for all proteins identified by mass spectrometry**

**Excel Table 3: List of high confidence lipid droplet proteins and their average iBAQ values**

**Excel Table 4: DHRS3 OE RNA-seq results, normalized counts**

**Excel Table 5: DHRS3 OE RNA-seq results, differentially expressed genes in DHRS3 OE vs. Control with padj < 0.05**

**Excel Table 6: RNA-seq results for indicated genes from the RNA-seq experiments depicted in Figure 7**

## References

1. Rambow, F., Marine, J.-C., and Goding, C.R. (2019). Melanoma plasticity and phenotypic diversity: therapeutic barriers and opportunities. Genes & Development 33, 1295–1318. 10.1101/gad.329771.119.

2. Rambow, F., Rogiers, A., Marin-Bejar, O., Aibar, S., Femel, J., Dewaele, M., Karras, P., Brown, D., Chang, Y.H., Debiec-Rychter, M., et al. (2018). Toward Minimal Residual Disease-Directed Therapy in Melanoma. Cell 174, 843–855.e819. 10.1016/j.cell.2018.06.025.

3. Hoek, K.S., Eichhoff, O.M., Schlegel, N.C., DolJbbeling, U., Kobert, N., Schaerer, L., Hemmi, S., and Dummer, R. (2008). In vivo Switching of Human Melanoma Cells between Proliferative and Invasive States. Cancer Research 68, 650–656. 10.1158/0008-5472.CAN-07-2491.

4. Tsoi, J., Robert, L., Paraiso, K., Galvan, C., Sheu, K.M., Lay, J., Wong, D.J.L., Atefi, M., Shirazi, R., Wang, X., et al. (2018). Multi-stage Differentiation Defines Melanoma Subtypes with Differential Vulnerability to Drug-Induced Iron-Dependent Oxidative Stress. Cancer Cell 33, 890–904.e895. 10.1016/j.ccell.2018.03.017.

5. Wouters, J., Kalender-Atak, Z., Minnoye, L., Spanier, K.I., De Waegeneer, M., Bravo González-Blas, C., Mauduit, D., Davie, K., Hulselmans, G., Najem, A., et al. (2020). Robust gene expression programs underlie recurrent cell states and phenotype switching in melanoma. Nature Cell Biology 22, 986–998. 10.1038/s41556-020-0547-3.

6. Karras, P., Bordeu, I., Pozniak, J., Nowosad, A., Pazzi, C., Van Raemdonck, N., Landeloos, E., Van Herck, Y., Pedri, D., Bervoets, G., et al. (2022). A cellular hierarchy in melanoma uncouples growth and metastasis. Nature 610, 190–198. 10.1038/s41586-022-05242-7.

7. Widmer, D.S., Cheng, P.F., Eichhoff, O.M., Belloni, B.C., Zipser, M.C., Schlegel, N.C., Javelaud, D., Mauviel, A., Dummer, R., and Hoek, K.S. (2012). Systematic classification of melanoma cells by phenotype-specific gene expression mapping. Pigment Cell & Melanoma Research 25, 343–353. 10.1111/j.1755-148X.2012.00986.x.

8. Ubellacker, J.M., Tasdogan, A., Ramesh, V., Shen, B., Mitchell, E.C., Martin-Sandoval, M.S., Gu, Z., McCormick, M.L., Durham, A.B., Spitz, D.R., et al. (2020). Lymph protects metastasizing melanoma cells from ferroptosis. Nature 585, 113–118. 10.1038/s41586-020-2623-z.

9. Pascual, G., Avgustinova, A., Mejetta, S., Martín, M., Castellanos, A., Attolini, C.S.-O., Berenguer, A., Prats, N., Toll, A., Hueto, J.A., et al. Targeting metastasis-initiating cells through the fatty acid receptor CD36.

10. Pascual, G., Avgustinova, A., Mejetta, S., Martín, M., Castellanos, A., Attolini, C.S.-O., Berenguer, A., Prats, N., Toll, A., Hueto, J.A., et al. (2017). Targeting metastasis-initiating cells through the fatty acid receptor CD36. Nature 541, 41–45. 10.1038/nature20791.

11. Broadfield, L.A., Duarte, J.A.G., Schmieder, R., Broekaert, D., Veys, K., Planque, M., Vriens, K., Karasawa, Y., Napolitano, F., Fujita, S., et al. (2021). Fat Induces Glucose Metabolism in Nontransformed Liver Cells and Promotes Liver Tumorigenesis. Cancer Research 81, 1988–2001. 10.1158/0008-5472.CAN-20-1954.

12. Lumaquin-Yin, D., Montal, E., Johns, E., Baggiolini, A., Huang, T.-H., Ma, Y., LaPlante, C., Suresh, S., Studer, L., and White, R.M. (2023). Lipid droplets are a metabolic vulnerability in melanoma. Nature Communications 14, 3192. 10.1038/s41467-023-38831-9.

13. Wilcock, D.J., Badrock, A.P., Wong, C.W., Owen, R., Guerin, M., Southam, A.D., Johnston, H., Telfer, B.A., Fullwood, P., Watson, J., et al. (2022). Oxidative stress from DGAT1 oncoprotein inhibition in melanoma suppresses tumor growth when ROS defenses are also breached. Cell Reports 39, 110995. 10.1016/j.celrep.2022.110995.

14. Zhang, M., Di Martino, J.S., Bowman, R.L., Campbell, N.R., Baksh, S.C., Simon-Vermot, T., Kim, I.S., Haldeman, P., Mondal, C., Yong-Gonzales, V., et al. (2018). Adipocyte-Derived Lipids Mediate Melanoma Progression via FATP Proteins. Cancer Discovery 8, 1006–1025. 10.1158/2159-8290.CD-17-1371.

15. Ting-Hsiang, H., Yilun, M., Emily, M., Shruthy, S., Mohita, M.T., Alexandra, C., Dianne, L., Nathaniel, R.C., Arianna, B., Richard, P.K., and Richard, M.W. (2022). Epigenetic reprogramming of melanoma cell state through fatty acid β-oxidation and Toll-like receptor 4 signaling. bioRxiv, 2022.2006.2016.496450. 10.1101/2022.06.16.496450.

16. Roberts, M.A., and Olzmann, J.A. (2020). Protein Quality Control and Lipid Droplet Metabolism. Annual Review of Cell and Developmental Biology 36, 115–139. 10.1146/annurev-cellbio-031320-101827.

17. Bersuker, K., and Olzmann, J.A. (2017). Establishing the lipid droplet proteome: Mechanisms of lipid droplet protein targeting and degradation. Biochimica et Biophysica Acta (BBA) - Molecular and Cell Biology of Lipids 1862, 1166–1177. 10.1016/j.bbalip.2017.06.006.

18. Krahmer, N., Guo, Y., Wilfling, F., Hilger, M., Lingrell, S., Heger, K., Newman, Heather W., Schmidt-Supprian, M., Vance, Dennis E., Mann, M., et al. (2011). Phosphatidylcholine Synthesis for Lipid Droplet Expansion Is Mediated by Localized Activation of CTP:Phosphocholine Cytidylyltransferase. Cell Metabolism 14, 504–515. 10.1016/j.cmet.2011.07.013.

19. Mejhert, N., Kuruvilla, L., Gabriel, K.R., Elliott, S.D., Guie, M.-A., Wang, H., Lai, Z.W., Lane, E.A., Christiano, R., Danial, N.N., et al. (2020). Partitioning of MLX-Family Transcription Factors to Lipid Droplets Regulates Metabolic Gene Expression. Molecular Cell 77, 1251–1264.e1259. 10.1016/j.molcel.2020.01.014.

20. Bersuker, K., Peterson, C.W.H., To, M., Sahl, S.J., Savikhin, V., Grossman, E.A., Nomura, D.K., and Olzmann, J.A. (2018). A Proximity Labeling Strategy Provides Insights into the Composition and Dynamics of Lipid Droplet Proteomes. Developmental Cell 44, 97–112.e117. 10.1016/j.devcel.2017.11.020.

21. Krahmer, N., Hilger, M., Kory, N., Wilfling, F., Stoehr, G., Mann, M., Farese, R.V., and Walther, T.C. (2013). Protein Correlation Profiles Identify Lipid Droplet Proteins with High Confidence*. Molecular & Cellular Proteomics 12, 1115–1126. 10.1074/mcp.M112.020230.

22. Krahmer, N., Najafi, B., Schueder, F., Quagliarini, F., Steger, M., Seitz, S., Kasper, R., Salinas, F., Cox, J., Uhlenhaut, N.H., et al. (2018). Organellar Proteomics and Phospho-Proteomics Reveal Subcellular Reorganization in Diet-Induced Hepatic Steatosis. Developmental Cell 47, 205–221.e207. 10.1016/j.devcel.2018.09.017.

23. Mejhert, N., Gabriel, K.R., Frendo-Cumbo, S., Krahmer, N., Song, J., Kuruvilla, L., Chitraju, C., Boland, S., Jang, D.-K., von Grotthuss, M., et al. (2022). The Lipid Droplet Knowledge Portal: A resource for systematic analyses of lipid droplet biology. Developmental Cell 57, 387–397.e384. 10.1016/j.devcel.2022.01.003.

24. Peterson, C.W.H., Deol, K.K., To, M., and Olzmann, J.A. (2021). Optimized protocol for the identification of lipid droplet proteomes using proximity labeling proteomics in cultured human cells. STAR Protocols 2, 100579. 10.1016/j.xpro.2021.100579.

25. Adams, M.K., Belyaeva, O.V., Wu, L., and Kedishvili, N.Y. (2014). The Retinaldehyde Reductase Activity of DHRS3 Is Reciprocally Activated by Retinol Dehydrogenase 10 to Control Retinoid Homeostasis*. Journal of Biological Chemistry 289, 14868–14880. 10.1074/jbc.M114.552257.

26. Fernandes-Silva, H., Araújo-Silva, H., Correia-Pinto, J., and Moura, R.S. (2020). Retinoic Acid: A Key Regulator of Lung Development. Biomolecules 10. 10.3390/biom10010152.

27. Inoue, Y., Hasegawa, S., Yamada, T., Date, Y., Mizutani, H., Nakata, S., Matsunaga, K., and Akamatsu, H. (2012). Bimodal effect of retinoic acid on melanocyte differentiation identified by time-dependent analysis. Pigment Cell & Melanoma Research 25, 299–311. 10.1111/j.1755-148X.2012.00988.x.

28. Roméro, C., Aberdam, E., Larnier, C., and Ortonne, J.-P. (1994). Retinoic acid as modulator of UVB-induced melanocyte differentiation: Involvement of the melanogenic enzymes expression. Journal of Cell Science 107, 1095–1103. 10.1242/jcs.107.4.1095.

29. Nusinow, D.P., Szpyt, J., Ghandi, M., Rose, C.M., McDonald, E.R., III, Kalocsay, M., Jané-Valbuena, J., Gelfand, E., Schweppe, D.K., Jedrychowski, M., et al. (2020). Quantitative Proteomics of the Cancer Cell Line Encyclopedia. Cell 180, 387–402.e316. 10.1016/j.cell.2019.12.023.

30. Deisenroth, C., Itahana, Y., Tollini, L., Jin, A., and Zhang, Y. (2011). p53-inducible DHRS3 Is an Endoplasmic Reticulum Protein Associated with Lipid Droplet Accumulation*. Journal of Biological Chemistry 286, 28343–28356. 10.1074/jbc.M111.254227.

31. Pataki, C.I., Rodrigues, J., Zhang, L., Qian, J., Efron, B., Hastie, T., Elias, J.E., Levitt, M., and Kopito, R.R. (2018). Proteomic analysis of monolayer-integrated proteins on lipid droplets identifies amphipathic interfacial alpha-helical membrane anchors. Proc Natl Acad Sci U S A 115, E8172–E8180. 10.1073/pnas.1807981115.

32. Louphrasitthiphol, P., Ledaki, I., Chauhan, J., Falletta, P., Siddaway, R., Buffa, F.M., Mole, D.R., Soga, T., and Goding, C.R. (2019). MITF controls the TCA cycle to modulate the melanoma hypoxia response. Pigment Cell & Melanoma Research 32, 792–808. 10.1111/pcmr.12802.

33. Vivas-García, Y., Falletta, P., Liebing, J., Louphrasitthiphol, P., Feng, Y., Chauhan, J., Scott, D.A., Glodde, N., Chocarro-Calvo, A., Bonham, S., et al. (2020). Lineage-Restricted Regulation of SCD and Fatty Acid Saturation by MITF Controls Melanoma Phenotypic Plasticity. Molecular Cell 77, 120–137.e129. 10.1016/j.molcel.2019.10.014.

34. Boncinelli, E., Simeone, A., Acampora, D., and Mavilio, F. (1991). HOX gene activation by retinoic acid. Trends in Genetics 7, 329–334. 10.1016/0168-9525(91)90423-N.

35. Breier, G., Bućan, M., Francke, U., Colberg_-_Poley, A.M., and Gruss, P. (1986). Sequential expression of murine homeo box genes during F9 EC cell differentiation. The EMBO Journal 5, 2209–2215-2215. 10.1002/j.1460-2075.1986.tb04486.x.

36. Tobin, R.P., Cogswell, D.T., Cates, V.M., Davis, D.M., Borgers, J.S.W., Van Gulick, R.J., Katsnelson, E., Couts, K.L., Jordan, K.R., Gao, D., et al. (2023). Targeting MDSC Differentiation Using ATRA: A Phase I/II Clinical Trial Combining Pembrolizumab and All-Trans Retinoic Acid for Metastatic Melanoma. Clinical Cancer Research 29, 1209–1219. 10.1158/1078-0432.CCR-22-2495.

37. Kane, M.A. (2012). Analysis, occurrence, and function of 9-cis-retinoic acid. Biochimica et Biophysica Acta (BBA) - Molecular and Cell Biology of Lipids 1821, 10–20. 10.1016/j.bbalip.2011.09.012.

38. Paterson, E.K., Ho, H., Kapadia, R., and Ganesan, A.K. (2013). 9-cis retinoic acid is the ALDH1A1 product that stimulates melanogenesis. Experimental Dermatology 22, 202–209. 10.1111/exd.12099.

39. Goding, C.R., and Arnheiter, H. (2019). MITF-the first 25 years. Genes Dev 33, 983–1007. 10.1101/gad.324657.119.

40. Mort, R.L., Jackson, I.J., and Patton, E.E. (2015). The melanocyte lineage in development and disease. Development 142, 620–632. 10.1242/dev.106567.

41. Louphrasitthiphol, P., Siddaway, R., Loffreda, A., Pogenberg, V., Friedrichsen, H., Schepsky, A., Zeng, Z., Lu, M., Strub, T., Freter, R., et al. (2020). Tuning Transcription Factor Availability through Acetylation-Mediated Genomic Redistribution. Molecular Cell 79, 472–487.e410. 10.1016/j.molcel.2020.05.025.

42. Aksan, I., and Goding, C.R. (1998). Targeting the Microphthalmia Basic Helix-Loop-Helix–Leucine Zipper Transcription Factor to a Subset of E-Box Elements In Vitro and In Vivo. Molecular and Cellular Biology 18, 6930–6938. 10.1128/MCB.18.12.6930.

43. Dorsky, R.I., Raible, D.W., and Moon, R.T. (2000). Direct regulation of nacre, a zebrafish MITF homolog required for pigment cell formation, by the Wnt pathway. Genes Dev 14, 158–162.

44. Takeda, K., Yasumoto, K.-i., Takada, R., Takada, S., Watanabe, K.-i., Udono, T., Saito, H., Takahashi, K., and Shibahara, S. (2000). Induction of Melanocyte-specific Microphthalmia-associated Transcription Factor by Wnt-3a*. Journal of Biological Chemistry 275, 14013–14016. 10.1074/jbc.C000113200.

45. Widlund, H.R., Horstmann, M.A., Price, E.R., Cui, J., Lessnick, S.L., Wu, M., He, X., and Fisher, D.E. (2002). β-Catenin–induced melanoma growth requires the downstream target Microphthalmia-associated transcription factor. Journal of Cell Biology 158, 1079–1087. 10.1083/jcb.200202049.

46. Schepsky, A., Bruser, K., Gunnarsson, G.J., Goodall, J., Hallsson, J.H., Goding, C.R., Steingrimsson, E., and Hecht, A. (2006). The Microphthalmia-Associated Transcription Factor Mitf Interacts with β-Catenin To Determine Target Gene Expression. Molecular and Cellular Biology 26, 8914–8927. 10.1128/MCB.02299-05.

47. Louphrasitthiphol, P., Chauhan, J., and Goding, C.R. (2020). ABCB5 is activated by MITF and β-catenin and is associated with melanoma differentiation. Pigment Cell & Melanoma Research 33, 112–118. 10.1111/pcmr.12830.

48. Chauhan, J.S., Hölzel, M., Lambert, J.-P., Buffa, F.M., and Goding, C.R. (2022). The MITF regulatory network in melanoma. Pigment Cell & Melanoma Research 35, 517–533. 10.1111/pcmr.13053.

49. Marusyk, A., Almendro, V., and Polyak, K. (2012). Intra-tumour heterogeneity: a looking glass for cancer? Nature Reviews Cancer 12, 323–334. 10.1038/nrc3261.

50. Gavish, A., Tyler, M., Greenwald, A.C., Hoefflin, R., Simkin, D., Tschernichovsky, R., Galili Darnell, N., Somech, E., Barbolin, C., Antman, T., et al. (2023). Hallmarks of transcriptional intratumour heterogeneity across a thousand tumours. Nature 618, 598–606. 10.1038/s41586-023-06130-4.

51. Hoek, K.S., and Goding, C.R. (2010). Cancer stem cells versus phenotype-switching in melanoma. Pigment Cell & Melanoma Research 23, 746–759. 10.1111/j.1755-148X.2010.00757.x.

52. Clement, E., Lazar, I., Attané, C., Carrié, L., Dauvillier, S., Ducoux-Petit, M., Esteve, D., Menneteau, T., Moutahir, M., Le Gonidec, S., et al. (2020). Adipocyte extracellular vesicles carry enzymes and fatty acids that stimulate mitochondrial metabolism and remodeling in tumor cells. The EMBO Journal 39, e102525. 10.15252/embj.2019102525.

53. Henderson, F., Johnston, H.R., Badrock, A.P., Jones, E.A., Forster, D., Nagaraju, R.T., Evangelou, C., Kamarashev, J., Green, M., Fairclough, M., et al. (2019). Enhanced Fatty Acid Scavenging and Glycerophospholipid Metabolism Accompany Melanocyte Neoplasia Progression in Zebrafish. Cancer Research 79, 2136–2151. 10.1158/0008-5472.CAN-18-2409.

54. Watt, M.J., Clark, A.K., Selth, L.A., Haynes, V.R., Lister, N., Rebello, R., Porter, L.H., Niranjan, B., Whitby, S.T., Lo, J., et al. (2019). Suppressing fatty acid uptake has therapeutic effects in preclinical models of prostate cancer. Science Translational Medicine 11, eaau5758. 10.1126/scitranslmed.aau5758.

55. Nieman, K.M., Kenny, H.A., Penicka, C.V., Ladanyi, A., Buell-Gutbrod, R., Zillhardt, M.R., Romero, I.L., Carey, M.S., Mills, G.B., Hotamisligil, G.S., et al. (2011). Adipocytes promote ovarian cancer metastasis and provide energy for rapid tumor growth. Nature Medicine 17, 1498–1503. 10.1038/nm.2492.

56. Lumaquin, D., Montal, E., Baggiolini, A., Ma, Y., LaPlante, C., Huang, T.-H., Suresh, S., Studer, L., and White, R.M. (2022). Lipid droplets are a metabolic vulnerability in melanoma. bioRxiv, 2022.2005.2004.490656. 10.1101/2022.05.04.490656.

57. Koundouros, N., and Poulogiannis, G. (2020). Reprogramming of fatty acid metabolism in cancer. British Journal of Cancer 122, 4–22. 10.1038/s41416-019-0650-z.

58. Park, Jun H., Vithayathil, S., Kumar, S., Sung, P.-L., Dobrolecki, Lacey E., Putluri, V., Bhat, Vadiraja B., Bhowmik, Salil K., Gupta, V., Arora, K., et al. (2016). Fatty Acid Oxidation-Driven Src Links Mitochondrial Energy Reprogramming and Oncogenic Properties in Triple-Negative Breast Cancer. Cell Reports 14, 2154–2165. 10.1016/j.celrep.2016.02.004.

59. Schlaepfer, I.R., Rider, L., Rodrigues, L.U., Gijón, M.A., Pac, C.T., Romero, L., Cimic, A., Sirintrapun, S.J., Glodé, L.M., Eckel, R.H., and Cramer, S.D. (2014). Lipid Catabolism via CPT1 as a Therapeutic Target for Prostate Cancer. Molecular Cancer Therapeutics 13, 2361–2371. 10.1158/1535-7163.MCT-14-0183.

60. Wang, T., Fahrmann, J.F., Lee, H., Li, Y.-J., Tripathi, S.C., Yue, C., Zhang, C., Lifshitz, V., Song, J., Yuan, Y., et al. (2018). JAK/STAT3-Regulated Fatty Acid β-Oxidation Is Critical for Breast Cancer Stem Cell Self-Renewal and Chemoresistance. Cell Metabolism 27, 136–150.e135. 10.1016/j.cmet.2017.11.001.

61. Doll, S., Proneth, B., Tyurina, Y.Y., Panzilius, E., Kobayashi, S., Ingold, I., Irmler, M., Beckers, J., Aichler, M., Walch, A., et al. (2017). ACSL4 dictates ferroptosis sensitivity by shaping cellular lipid composition. Nature Chemical Biology 13, 91–98. 10.1038/nchembio.2239.

62. Magtanong, L., Ko, P.-J., To, M., Cao, J.Y., Forcina, G.C., Tarangelo, A., Ward, C.C., Cho, K., Patti, G.J., Nomura, D.K., et al. (2019). Exogenous Monounsaturated Fatty Acids Promote a Ferroptosis-Resistant Cell State. Cell Chemical Biology 26, 420–432.e429. 10.1016/j.chembiol.2018.11.016.

63. Dierge, E., Debock, E., Guilbaud, C., Corbet, C., Mignolet, E., Mignard, L., Bastien, E., Dessy, C., Larondelle, Y., and Feron, O. (2021). Peroxidation of n-3 and n-6 polyunsaturated fatty acids in the acidic tumor environment leads to ferroptosis-mediated anticancer effects. Cell Metabolism 33, 1701–1715.e1705. 10.1016/j.cmet.2021.05.016.

64. Mathiowetz, A.J., and Olzmann, J.A. (2024). Lipid droplets and cellular lipid flux. Nature Cell Biology. 10.1038/s41556-024-01364-4.

65. Cruz, A.L.S., Barreto, E.d.A., Fazolini, N.P.B., Viola, J.P.B., and Bozza, P.T. (2020). Lipid droplets: platforms with multiple functions in cancer hallmarks. Cell Death & Disease 11, 105. 10.1038/s41419-020-2297-3.

66. Herker, E., Vieyres, G., Beller, M., Krahmer, N., and Bohnert, M. (2021). Lipid Droplet Contact Sites in Health and Disease. Trends in Cell Biology 31, 345–358. 10.1016/j.tcb.2021.01.004.

67. Perera, R.M., Stoykova, S., Nicolay, B.N., Ross, K.N., Fitamant, J., Boukhali, M., Lengrand, J., Deshpande, V., Selig, M.K., Ferrone, C.R., et al. (2015). Transcriptional control of autophagy–lysosome function drives pancreatic cancer metabolism. Nature 524, 361–365. 10.1038/nature14587.

68. Yang, S., Wang, X., Contino, G., Liesa, M., Sahin, E., Ying, H., Bause, A., Li, Y., Stommel, J.M., Dell’Antonio, G., et al. (2011). Pancreatic cancers require autophagy for tumor growth. Genes & Development 25, 717–729.

69. Audet-Walsh, É., Vernier, M., Yee, T., Laflamme, C., Li, S., Chen, Y., and Giguère, V. (2018). SREBF1 Activity Is Regulated by an AR/mTOR Nuclear Axis in Prostate Cancer. Molecular Cancer Research 16, 1396–1405. 10.1158/1541-7786.MCR-17-0410.

70. Chauhan, S.S., Casillas, A.L., Vizzerra, A.D., Liou, H., Clements, A.N., Flores, C.E., Prevost, C.T., Kashatus, D.F., Snider, A.J., Snider, J.M., and Warfel, N.A. (2024). PIM1 drives lipid droplet accumulation to promote proliferation and survival in prostate cancer. Oncogene 43, 406–419. 10.1038/s41388-023-02914-0.

71. Lee, G., Zheng, Y., Cho, S., Jang, C., England, C., Dempsey, J.M., Yu, Y., Liu, X., He, L., Cavaliere, P.M., et al. (2017). Post-transcriptional Regulation of De Novo Lipogenesis by mTORC1-S6K1-SRPK2 Signaling. Cell 171, 1545–1558.e1518. 10.1016/j.cell.2017.10.037.

72. Guri, Y., Colombi, M., Dazert, E., Hindupur, S.K., Roszik, J., Moes, S., Jenoe, P., Heim, M.H., Riezman, I., Riezman, H., and Hall, M.N. (2017). mTORC2 Promotes Tumorigenesis via Lipid Synthesis. Cancer Cell 32, 807–823.e812. 10.1016/j.ccell.2017.11.011.

73. Accioly, M.T., Pacheco, P., Maya-Monteiro, C.M., Carrossini, N., Robbs, B.K., Oliveira, S.S., Kaufmann, C., Morgado-Diaz, J.A., Bozza, P.T., and Viola, J.o.P.B. (2008). Lipid Bodies Are Reservoirs of Cyclooxygenase-2 and Sites of Prostaglandin-E2 Synthesis in Colon Cancer Cells. Cancer Research 68, 1732–1740. 10.1158/0008-5472.CAN-07-1999.

74. Tirinato, L., Liberale, C., Di Franco, S., Candeloro, P., Benfante, A., La Rocca, R., Potze, L., Marotta, R., Ruffilli, R., Rajamanickam, V.P., et al. (2015). Lipid Droplets: A New Player in Colorectal Cancer Stem Cells Unveiled by Spectroscopic Imaging. Stem Cells 33, 35–44. 10.1002/stem.1837.

75. Li, J., Condello, S., Thomes-Pepin, J., Ma, X., Xia, Y., Hurley, T.D., Matei, D., and Cheng, J.-X. (2017). Lipid Desaturation Is a Metabolic Marker and Therapeutic Target of Ovarian Cancer Stem Cells. Cell Stem Cell 20, 303–314.e305. 10.1016/j.stem.2016.11.004.

76. Kostecka, L.G., Mendez, S., Li, M., Khare, P., Zhang, C., Le, A., Amend, S.R., and Pienta, K.J. (2024). Cancer cells employ lipid droplets to survive toxic stress. The Prostate n/a. 10.1002/pros.24680.

77. Nardi, F., Franco, O.E., Fitchev, P., Morales, A., Vickman, R.E., Hayward, S.W., and Crawford, S.E. (2019). DGAT1 Inhibitor Suppresses Prostate Tumor Growth and Migration by Regulating Intracellular Lipids and Non-Centrosomal MTOC Protein GM130. Scientific Reports 9, 3035. 10.1038/s41598-019-39537-z.

78. Wang, Y., Hu, Y., Xu, R., Jin, X., and Jiao, W. (2024). Plin2 inhibits autophagy via activating AKT/mTOR pathway in non-small cell lung cancer. Experimental Cell Research 435, 113955. 10.1016/j.yexcr.2024.113955.

79. Roberts, M.A., Deol, K.K., Mathiowetz, A.J., Lange, M., Leto, D.E., Stevenson, J., Hashemi, S.H., Morgens, D.W., Easter, E., Heydari, K., et al. (2023). Parallel CRISPR-Cas9 screens identify mechanisms of PLIN2 and lipid droplet regulation. Developmental Cell 58, 1782–1800.e1710. 10.1016/j.devcel.2023.07.001.

80. Liu, W., Liu, X., Liu, Y., Ling, T., Chen, D., Otkur, W., Zhao, H., Ma, M., Ma, K., Dong, B., et al. (2022). PLIN2 promotes HCC cells proliferation by inhibiting the degradation of HIF1α. Experimental Cell Research 418, 113244. 10.1016/j.yexcr.2022.113244.

81. Carreira, S., Goodall, J., Aksan, I., La Rocca, S.A., Galibert, M.-D., Denat, L., Larue, L., and Goding, C.R. (2005). Mitf cooperates with Rb1 and activates p21Cip1 expression to regulate cell cycle progression. Nature 433, 764–769. 10.1038/nature03269.

82. Vlčková, K., Vachtenheim, J., Réda, J., Horák, P., and Ondrušová, L. (2018). Inducibly decreased MITF levels do not affect proliferation and phenotype switching but reduce differentiation of melanoma cells. Journal of Cellular and Molecular Medicine 22, 2240–2251. 10.1111/jcmm.13506.

83. Zhang, P., Liu, W., Zhu, C., Yuan, X., Li, D., Gu, W., Ma, H., Xie, X., and Gao, T. (2012). Silencing of GPNMB by siRNA Inhibits the Formation of Melanosomes in Melanocytes in a MITF-Independent Fashion. PLOS ONE 7, e42955. 10.1371/journal.pone.0042955.

84. Kolberg, L., Raudvere, U., Kuzmin, I., Adler, P., Vilo, J., and Peterson, H. (2023). g:Profiler—interoperable web service for functional enrichment analysis and gene identifier mapping (2023 update). Nucleic Acids Research 51, W207–W212. 10.1093/nar/gkad347.

85. Cox, J., and Mann, M. (2008). MaxQuant enables high peptide identification rates, individualized p.p.b.-range mass accuracies and proteome-wide protein quantification. Nature Biotechnology 26, 1367–1372. 10.1038/nbt.1511.

86. Tyanova, S., Temu, T., and Cox, J. (2016). The MaxQuant computational platform for mass spectrometry-based shotgun proteomics. Nature Protocols 11, 2301–2319. 10.1038/nprot.2016.136.

87. Tyanova, S., Temu, T., Sinitcyn, P., Carlson, A., Hein, M.Y., Geiger, T., Mann, M., and Cox, J. (2016). The Perseus computational platform for comprehensive analysis of (prote)omics data. Nature Methods 13, 731–740. 10.1038/nmeth.3901.

88. Patro, R., Duggal, G., Love, M.I., Irizarry, R.A., and Kingsford, C. (2017). Salmon provides fast and bias-aware quantification of transcript expression. Nature Methods 14, 417–419. 10.1038/nmeth.4197.

89. Bolger, A.M., Lohse, M., and Usadel, B. (2014). Trimmomatic: a flexible trimmer for Illumina sequence data. Bioinformatics 30, 2114–2120. 10.1093/bioinformatics/btu170.

90. Love, M.I., Huber, W., and Anders, S. (2014). Moderated estimation of fold change and dispersion for RNA-seq data with DESeq2. Genome Biology 15, 550. 10.1186/s13059-014-0550-8.

91. Robinson, M.D., McCarthy, D.J., and Smyth, G.K. (2010). edgeR: a Bioconductor package for differential expression analysis of digital gene expression data. Bioinformatics 26, 139–140. 10.1093/bioinformatics/btp616.

92. Dobin, A., Davis, C.A., Schlesinger, F., Drenkow, J., Zaleski, C., Jha, S., Batut, P., Chaisson, M., and Gingeras, T.R. (2013). STAR: ultrafast universal RNA-seq aligner. Bioinformatics 29, 15–21. 10.1093/bioinformatics/bts635.

93. Martin, M. (2011). Cutadapt removes adapter sequences from high-throughput sequencing reads. EMBnet.journal; Vol 17, No 1: Next Generation Sequencing Data Analysis. 10.14806/ej.17.1.200.

94. Heinz, S., Benner, C., Spann, N., Bertolino, E., Lin, Y.C., Laslo, P., Cheng, J.X., Murre, C., Singh, H., and Glass, C.K. (2010). Simple Combinations of Lineage-Determining Transcription Factors Prime cis-Regulatory Elements Required for Macrophage and B Cell Identities. Molecular Cell 38, 576–589. 10.1016/j.molcel.2010.05.004.

95. Gennady, K., Vladimir, S., Nikolay, B., Boris, S., Maxim, N.A., and Alexey, S. (2021). Fast gene set enrichment analysis. bioRxiv, 060012. 10.1101/060012.

96. Rappsilber, J., Mann, M., and Ishihama, Y. (2007). Protocol for micro-purification, enrichment, pre-fractionation and storage of peptides for proteomics using StageTips. Nature Protocols 2, 1896–1906. 10.1038/nprot.2007.261.

97. Olsen, J.V., de Godoy, L.M.F., Li, G., Macek, B., Mortensen, P., Pesch, R., Makarov, A., Lange, O., Horning, S., and Mann, M. (2005). Parts per Million Mass Accuracy on an Orbitrap Mass Spectrometer via Lock Mass Injection into a C-trap. Molecular & Cellular Proteomics 4, 2010–2021. 10.1074/mcp.T500030-MCP200.

98. Bunkenborg, J., García, G.E., Paz, M.I.P., Andersen, J.S., and Molina, H. (2010). The minotaur proteome: Avoiding cross-species identifications deriving from bovine serum in cell culture models. PROTEOMICS 10, 3040–3044. 10.1002/pmic.201000103.

99. Schwanhäusser, B., Busse, D., Li, N., Dittmar, G., Schuchhardt, J., Wolf, J., Chen, W., and Selbach, M. (2011). Global quantification of mammalian gene expression control. Nature 473, 337–342. 10.1038/nature10098.

100. Hao, Y., Stuart, T., Kowalski, M.H., Choudhary, S., Hoffman, P., Hartman, A., Srivastava, A., Molla, G., Madad, S., Fernandez-Granda, C., and Satija, R. (2024). Dictionary learning for integrative, multimodal and scalable single-cell analysis. Nature Biotechnology 42, 293–304. 10.1038/s41587-023-01767-y.

101. Hafemeister, C., and Satija, R. (2019). Normalization and variance stabilization of single-cell RNA-seq data using regularized negative binomial regression. Genome Biology 20, 296. 10.1186/s13059-019-1874-1.

102. Subramanian, A., Tamayo, P., Mootha, V.K., Mukherjee, S., Ebert, B.L., Gillette, M.A., Paulovich, A., Pomeroy, S.L., Golub, T.R., Lander, E.S., and Mesirov, J.P. (2005). Gene set enrichment analysis: a knowledge-based approach for interpreting genome-wide expression profiles. Proc Natl Acad Sci U S A 102, 15545–15550.

