## Supplementary Figures for "The lipid droplet protein DHRS3 is a regulator of melanoma cell state"

### Supplementary Figure 1

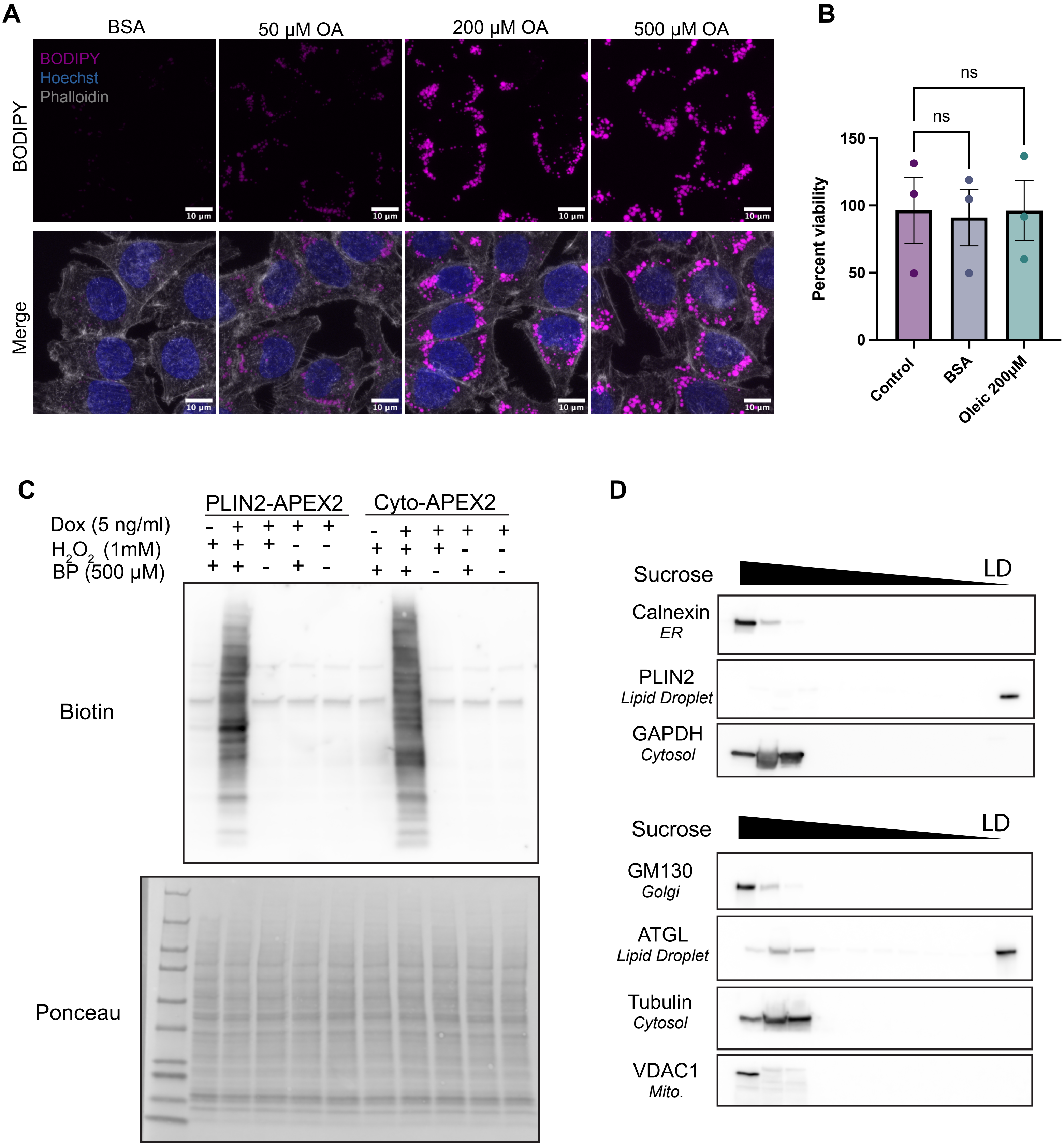

### Supplementary Figure 3

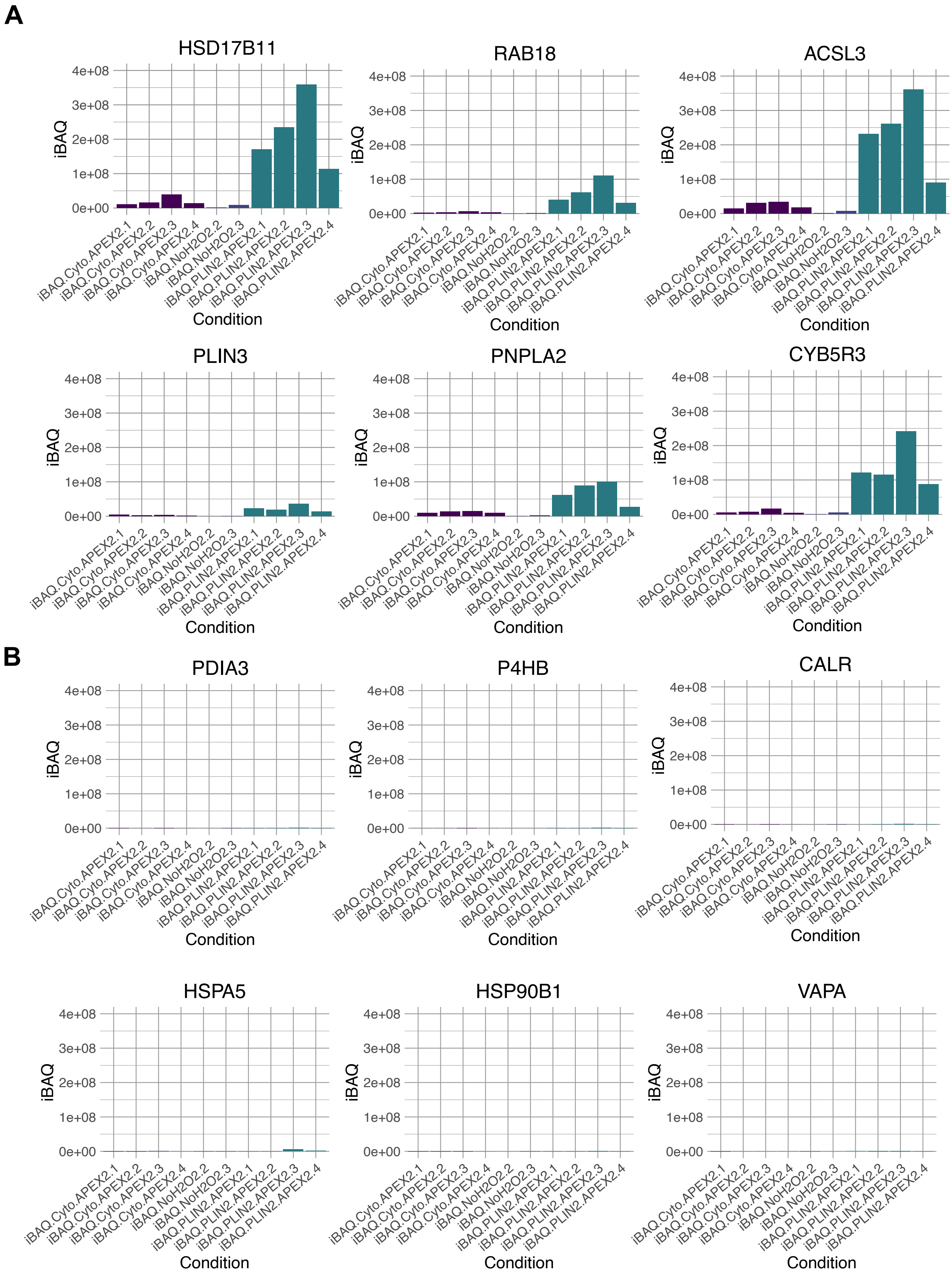

### Supplementary Figure 2

**A**

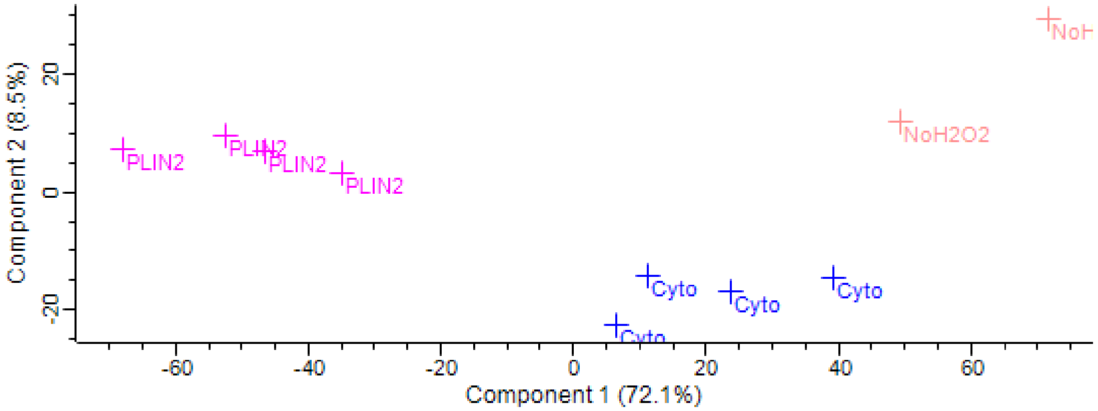

### Supplementary Figure 4

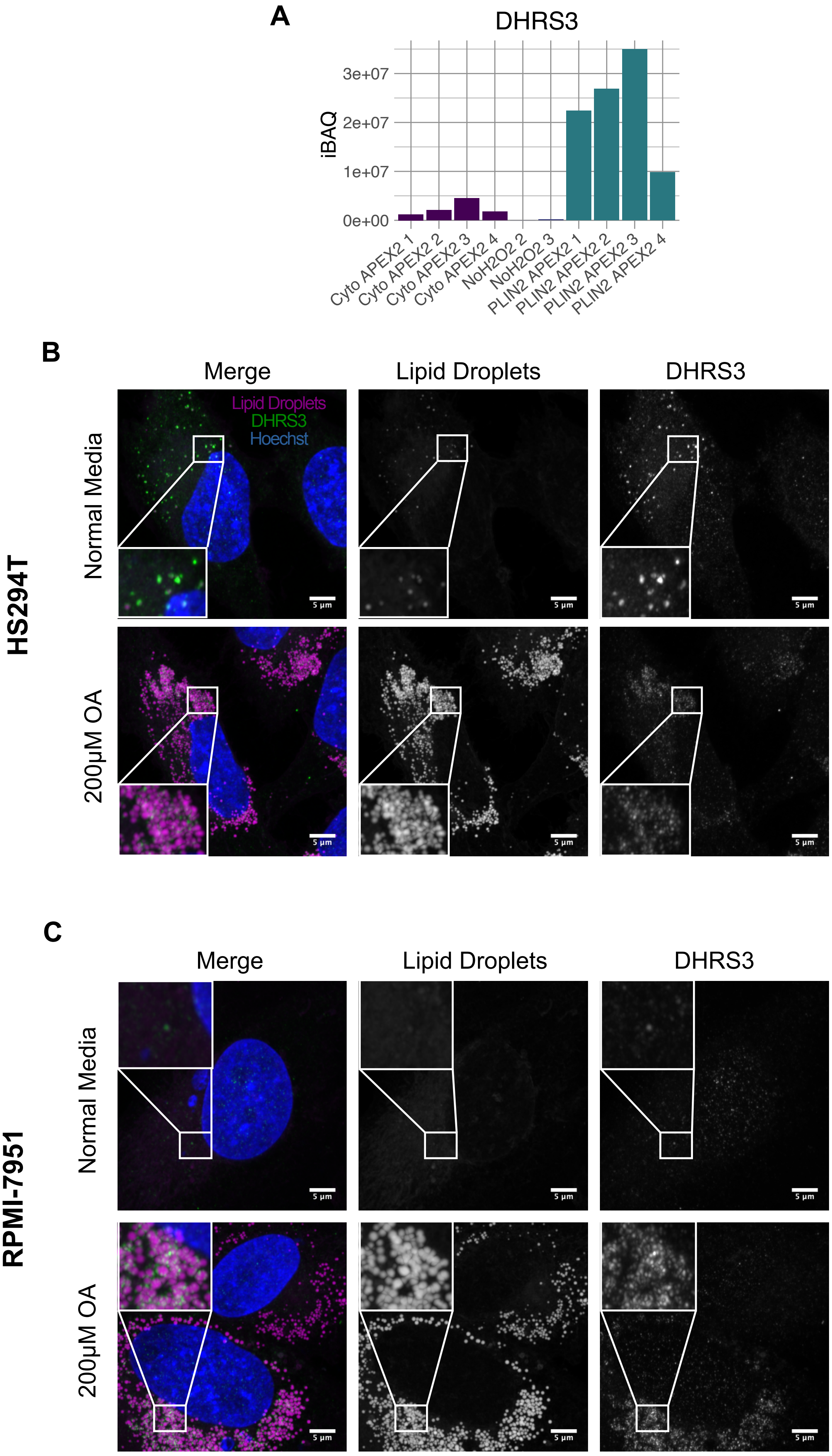

### Supplementary Figure 5

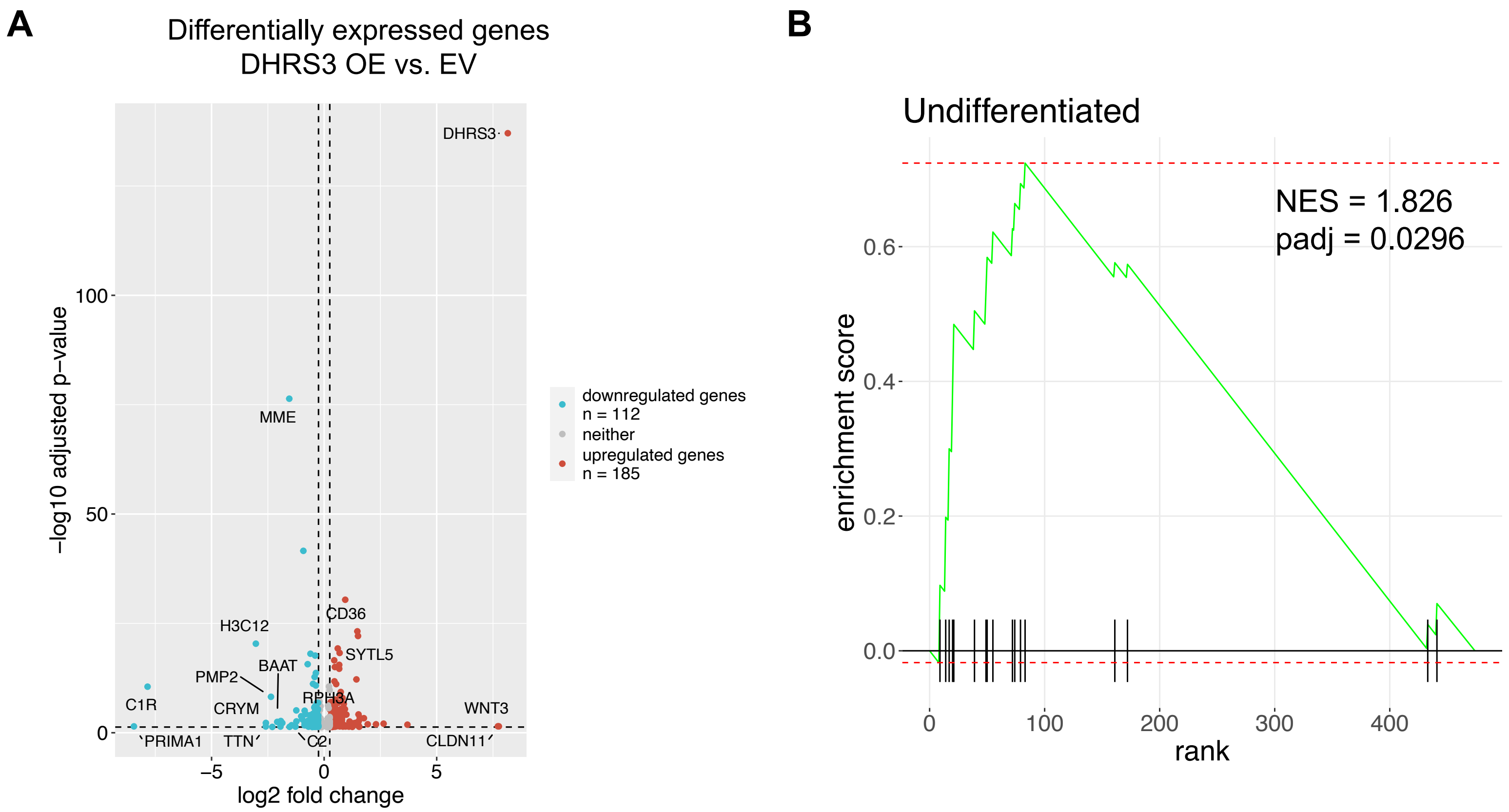


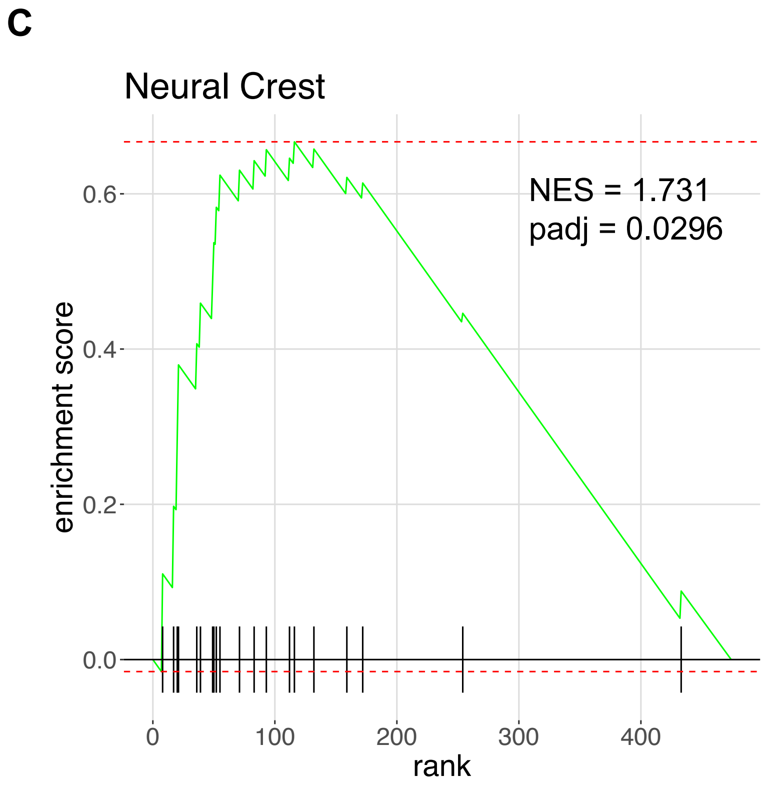


**D**

**Enriched Motifs with DHRS3 Overexpression**

| Rank | Motif | Best match | p-value | q-value |
|---|---|---|---|---|
| **1** | 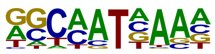 | *Hoxa9* | 1e-4 | 0.0434 |
| **2** | 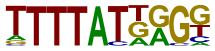 | *Hoxb13* | 1e-3 | 0.0434 |
| **3** | 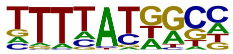 | *Hoxa11* | 1e-3 | 0.0434 |
| **4** | 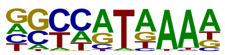 | *Hoxb11* | 1e-3 | 0.0618 |

**Supplementary Figure 1. Validation of lipid droplet isolation with APEX2 and gradient ultracentrifugation. Related to Figure 1.** A. Titration of Oleic Acid in A375 cells for 24 hours followed by fixation and imaging. Scale bars are 5 μm. B. Cyquant to measure cell viability after 24-hour treatment with 200 μM Oleic Acid. Data are normalized to the average of the control. Significance measured with One way ANOVA with Dunnett's Multiple Hypothesis Correction. Data are presented as mean +/- SD. C. Western blot of Cyto-V5-APEX2 and PLIN2-V5-APEX2 cell lines treated +/- $H_2O_2$ for 1 min, +/- Biotin Phenol for 30 min, and Doxycycline for 48 hours at the indicated doses D. Representative western blot results for fractionation of lipid droplets from wild-type A375 cells (following 24-hour treatment with Oleic Acid). Fractions were loaded in equal percent by volume. Data in A-D are representative of 3 independent experiments.

**Supplementary Figure 2. Analysis of LC MS/MS results. Related to Figure 2.** PCA plot of the 10 conditions submitted for LC-MS/MS. Each condition clusters separately indicating true differences between the proteins identified with each.

**Supplementary Figure 3. Quality control of lipid droplet proteomic data. Related to Figure 2**. A-B. iBAQ values of commonly identified lipid droplets (A) and common contaminants (B).

**Supplementary Figure 4. Localization of DHRS3 to the lipid droplet in undifferentiated melanoma cells. Related to Figure 4.** A. iBAQ values from the lipid droplet proteomics dataset for DHRS3. B-C Representative images of HS294T (B) and RPMI-7951 (C) stained for lipid droplets (LipidTOX) and endogenous DHRS3 in normal media and media with 200 μM Oleic Acid for 24 hours. Scale bars are 5 μm.

**Supplementary Figure 5. Differential gene expression & pathway analysis of DHRS3 RNA-seq. Related to Figure 5 and 6.** A. Volcano plot showing differentially expressed genes in DHRS3 OE vs. Control cells. B-C. Enrichment plots of Tsoi undifferentiated (B) and neural crest-like (C) pathways in DHRS3 OE cells. D. Top known motifs from HOMER motif enrichment analysis of the upregulated genes in DHRS3 OE cells (with $p<0.05$).
